# Video-evoked fMRI BOLD responses are highly consistent across different data acquisition sites

**DOI:** 10.1101/2021.10.04.463088

**Authors:** Lisa Byrge, Dorit Kliemann, Ye He, Hu Cheng, J. Michael Tyszka, Ralph Adolphs, Daniel P. Kennedy

## Abstract

Naturalistic imaging paradigms, in which participants view complex videos in the scanner, are increasingly used in human cognitive neuroscience. Videos evoke temporally synchronized brain responses that are similar across subjects as well as within subjects, but the reproducibility of these brain responses across different data acquisition sites has not yet been quantified. Here we characterize the consistency of brain responses across independent samples of participants viewing the same videos in fMRI scanners at different sites (Indiana University and Caltech). We compared brain responses collected at these different sites for two carefully matched datasets with identical scanner models, acquisition, and preprocessing details, along with a third unmatched dataset in which these details varied. Our overall conclusion is that for matched and unmatched datasets alike, video-evoked brain responses have high consistency across these different sites, both when compared across groups and across pairs of individuals. As one might expect, differences between sites were larger for unmatched datasets than matched datasets. Residual differences between datasets could in part reflect participant-level variability rather than scanner- or data-related effects. Altogether our results indicate promise for the development and, critically, generalization of video fMRI studies of individual differences in healthy and clinical populations alike.

## Introduction

Problems with reproducibility and reliability of scientific findings have arisen across numerous fields over the past two decades (Ioannidis, 2005). Functional magnetic resonance imaging (fMRI) studies have been far from immune, with inconsistent results found across numerous fMRI paradigms (Nickerson, 2018; Poldrack et al., 2017; Zuo et al., 2019; He et al., 2020; Elliott et al., 2020). Inconsistent results could indicate true and potentially relevant differences in study populations. But different data processing and data analysis choices can yield different conclusions from the same datasets (e.g., Eklund et al., 2016; Botvinik-Nezer, et al., 2020), and datasets collected from different scanners at different sites can contain non-biological variability due to differences in scanners and protocols (Friedman et al, 2006; Yu et al., 2018). Altogether these considerations indicate the importance of directly testing reproducibility across datasets collected at different sites.

Naturalistic viewing fMRI, or video fMRI (here; vfMRI; Hasson et al., 2004), has emerged in recent years as an attractive alternative to conventional task- and connectivity-based paradigms. Videos are arguably more ecologically valid, and permit greater compliance in the scanner (Eickhoff, Milham, & Vanderwal, 2020; Vanderwal, Eilbott, & Castellanos, 2019) making them an ideal candidate to use for developmental and clinical samples (e.g., Richardson et al., 2019). While vfMRI data can be analyzed using conventional task- and connectivity-based approaches, a distinct analysis approach that is based on measuring similarity or synchrony among participant brain responses has gained prominence (Saarimäki, 2021). This inter-subject correlation-based approach (ISC; Hasson et al., 2004) presents its own distinct analytic requirements due to the dependencies inherent in similarity measurements (Chen et al., 2017; Nastase et al., 2019). Within the same dataset from the same scanner, vfMRI paradigms can evoke markedly similar responses across subjects in many parts of the brain (e.g., Hasson et al., 2004, 2009, 2010; Byrge, Dubois, et al., 2015; Richardson et al., 2018; Nastase et al., 2019). Video-evoked brain responses have also been shown to be reliable within individual subjects after repeated stimulus presentations, in some regions (for review, see Hasson et al., 2010). Reliable responses are observed most consistently throughout posterior swaths of cortex including visual and auditory primary sensory and association areas and, for some video stimuli, can also extend to include parts of default network and lateral prefrontal cortex (Hasson et al., 2009, 2010; Byrge, Dubois, et al., 2015; Burunat et al., 2016). However, the extent to which brain responses during vfMRI are reproducible across different datasets collected at different sites has not yet been examined.

This issue of examining reproducibility of vfMRI across different sites takes on increased importance given recent momentum toward using vfMRI for clinical studies (autism: Hasson et al., 2009, Salmi et al., 2013, Byrge, Dubois, et al., 2015; schizophrenia: Yang et al., 2020; depression: Guo et al., 2015, Gruskin et al., 2020). The idea is to first use vfMRI to establish “normative” or “benchmark” patterns of brain responses to a video stimulus with clinically relevant features. This then makes it possible to quantify the extent to which an individual’s brain responses deviate from this reference pattern, in some particular brain area(s) or at some particular moment(s) of the video (Hasson et al., 2010; Eickhoff et al., 2020). The hope is that the combination of rich, dynamic stimuli that engage multiple brain networks simultaneously, the relative ease of standardizing stimuli and protocols across different data sites, and the increased data quantity and quality permitted by greater scan compliance might yield insights into the neural basis for the given condition, facilitate discovery of novel biomarkers (Sonkusare et al., 2019; Eickhoff et al., 2020), and eventually inform diagnosis as well as measure efficacy of interventions (Hasson et al., 2010).

Many clinical neuroscience studies are moving to multi-site consortiums (e.g., Di Martino et al., 2017, Loth et al., 2017), which aggregate data collection across different sites to obtain an appropriate sample. However, the weak point in clinical neuroscience studies can often be generalization of findings across different studies, samples, and sites (e.g., Kliemann et al., 2018, King et al., 2019; He et al., 2020). This presumably occurs due to combinations of factors that can include individual variability, methodological and stimulus variation, and differences between scanner equipment and standardization. Using video stimuli can minimize methodological and stimulus variation, as noted. But there is considerable individual variation in brain organization and function within the healthy “control” population (Zilles & Amunts, 2013; Dubois & Adolphs, 2016; Holmes & Patrick, 2018), including trait-linked variation in video-evoked brain response similarity (e.g., Salmi et al., 2013; Finn et al., 2018). It is therefore important to test the extent to which the “normative” pattern of brain responding to a video is itself reproducible across different sites, before using it as a clinical reference or benchmark. Such an investigation may also provide insights for fMRI harmonization efforts more generally. This is because stimulus-driven brain responses permit partitioning of variance between exogenously- and endogenously-driven brain function in a way that is not possible for some other types of widely-used fMRI paradigms like resting state functional connectivity.

Thus, here we directly examine cross-site consistency of evoked brain responses during video scans collected at two different data sites, Indiana University (Indiana) and California Institute of Technology (Caltech) in independent samples of healthy adults. The primary datasets for this manuscript are carefully matched datasets that were collected on different physical scanners in different states, but with potential sources of cross-site variability tightly controlled: identical scanner models, identical scan protocols, identical preprocessing pipelines, and identical analysis procedures (Table 1). Characterizing the similarity of brain responses across these closely matched datasets in a sample of typical controls will thus suggest a potential upper bound on the levels of cross-site consistency to be expected when the same video stimuli are used and other details are matched as closely as possible. As a further exploratory step, we also examined cross-site similarity of brain responses between two unmatched datasets: the Caltech dataset and an earlier pilot dataset (Pilot) also collected at Indiana University, but several years earlier and prior to a scanner upgrade. This Pilot dataset uses the same video stimuli, but different scanner models, different scan protocols, and differences across numerous dimensions of preprocessing approaches (Table 1). Although the unmatched acquisitions were not designed to disentangle specific sources of cross-site variability, we include that comparison as a case study that is informative about the ranges of similarity possible when sources of cross-site variation vary somewhat more freely – as is the case in some multi-site studies, particularly those pooled from pre-existing datasets. Thus, here we map out where in the brain to expect more consistent responses across sites, and conversely, where variability across matched datasets and unmatched datasets most strongly manifests in vfMRI paradigms. This establishes a key foundation for the clinical use of vfMRI, because confidently identifying atypical video-evoked responses in particular brain regions is ultimately limited by the reliability of vfMRI in that region (see also Elliott et al., 2021).

**Table 1.**
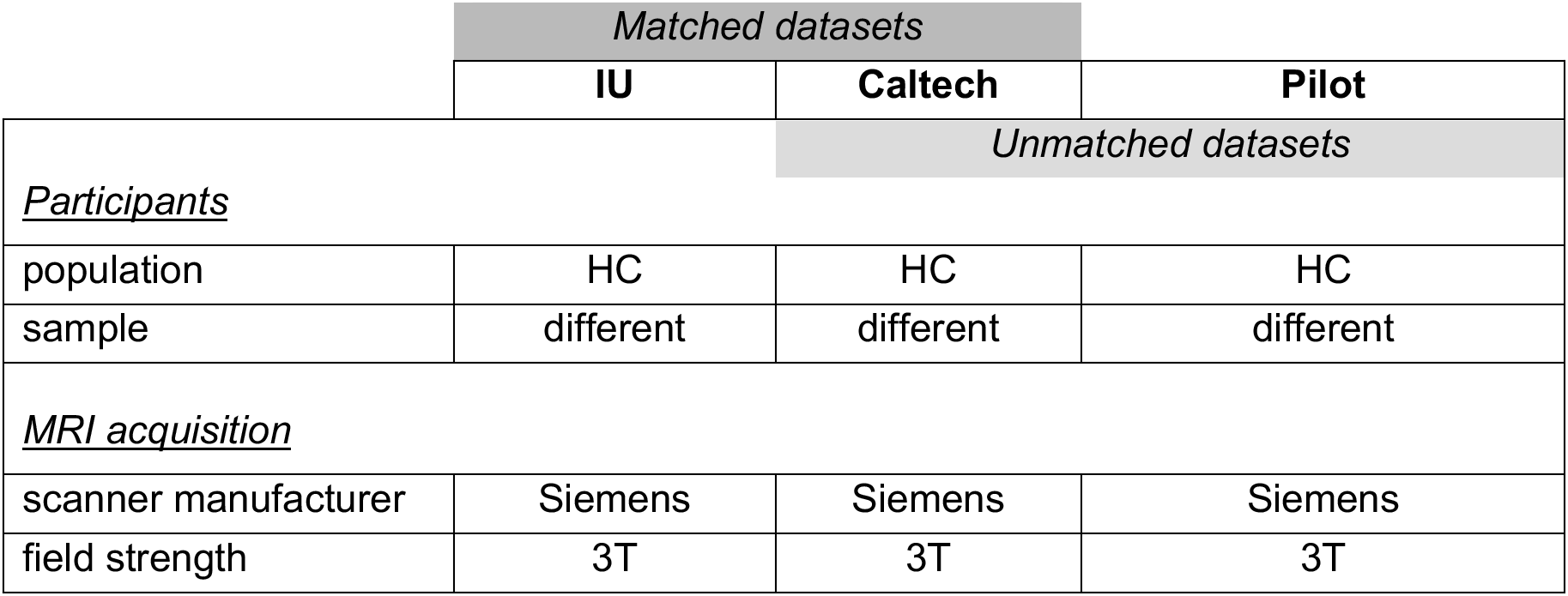
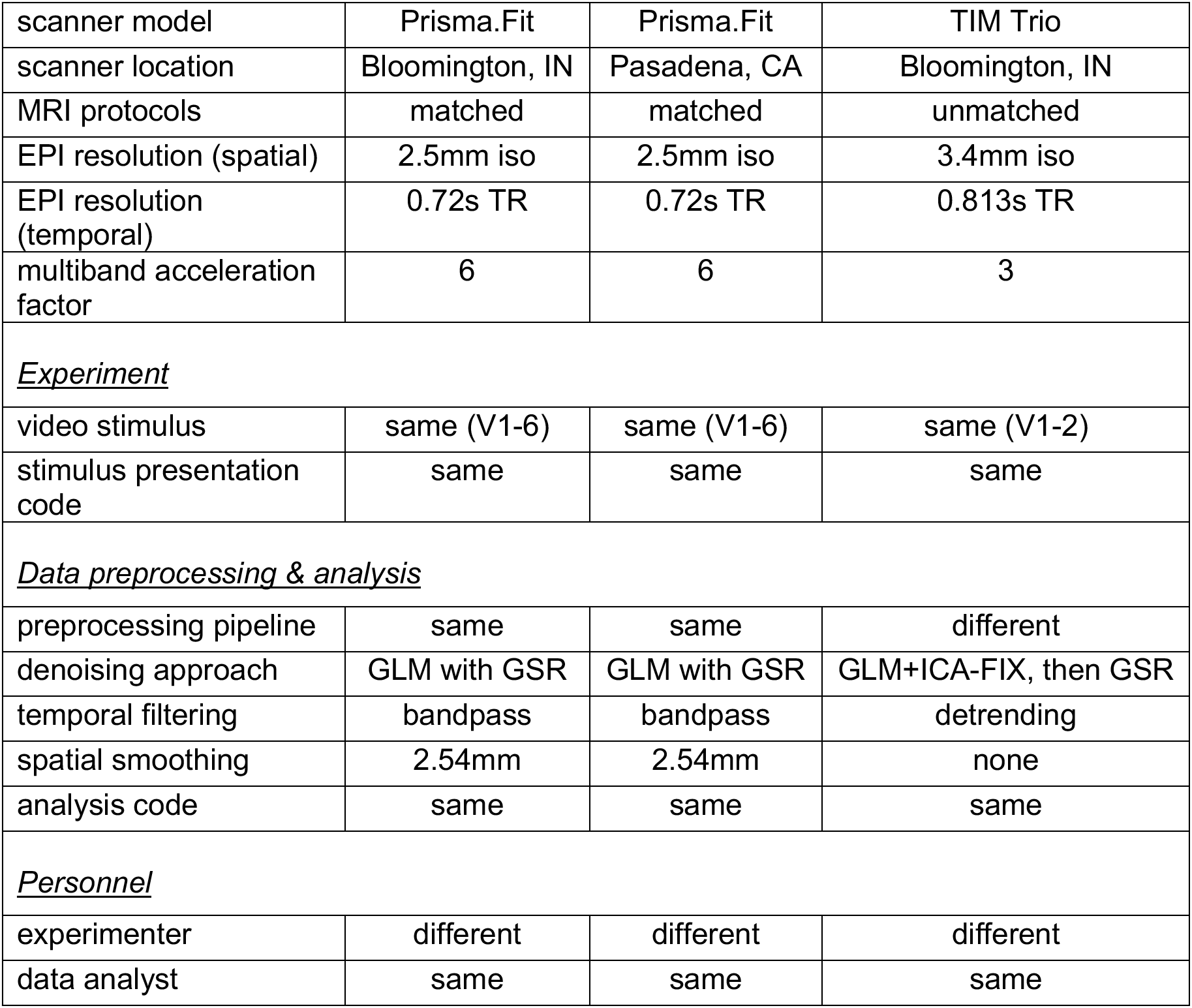
Similarities and differences between the matched and unmatched datasets. Table presents the main similarities and differences between the matched (IU & Caltech; dark gray) and unmatched (Caltech & Pilot; light gray) datasets. Table organization corresponds roughly to the taxonomy of reproducibility in neuroimaging from Nichols and colleagues (2017). The primary comparison between matched datasets is situated between “Near replicability” and “Intermediate replicability” of generalization over materials and methods in that taxonomy. The exploratory comparison between unmatched datasets is situated between “Intermediate replicability” and “Far replicability”; for that comparison, the Pilot acquisition was resampled temporally to match the sampling rate of the primary matched datasets. HC, healthy control adults. No participants overlapped between datasets. Entries listed as “same” and “different” for brevity are further detailed in Methods.

|  | Matched datasets |  |  |
| --- | --- | --- | --- |
|  | IU | Caltech | Pilot |
| <u>Participants</u> |  | Unmatched datasets |  |
| population | HC | HC | HC |
| sample | different | different | different |
| <u>MRI acquisition</u> |  |  |  |
| scanner manufacturer | Siemens | Siemens | Siemens |
| field strength | 3T | 3T | 3T |
| scanner model | Prisma.Fit | Prisma.Fit | TIM Trio |
| scanner location | Bloomington, IN | Pasadena, CA | Bloomington, IN |
| MRI protocols | matched | matched | unmatched |
| EPI resolution (spatial) | 2.5mm iso | 2.5mm iso | 3.4mm iso |
| EPI resolution (temporal) | 0.72s TR | 0.72s TR | 0.813s TR |
| multiband acceleration factor | 6 | 6 | 3 |
| <u>Experiment</u> |  |  |  |
| video stimulus | same (V1-6) | same (V1-6) | same (V1-2) |
| stimulus presentation code | same | same | same |
| <u>Data preprocessing & analysis</u> |  |  |  |
| preprocessing pipeline | same | same | different |
| denoising approach | GLM with GSR | GLM with GSR | GLM+ICA-FIX, then GSR |
| temporal filtering | bandpass | bandpass | detrending |
| spatial smoothing | 2.54mm | 2.54mm | none |
| analysis code | same | same | same |
| <u>Personnel</u> |  |  |  |
| experimenter | different | different | different |
| data analyst | same | same | same |

## Materials & Methods

### Participants

#### Matched datasets

The primary matched datasets were collected at two sites, Indiana University and Caltech, between 2017 and 2020, as part of a larger project including both typically developed adults and adults with autism spectrum disorder (ASD). Only data from typically developed adults are included in the current report (*N* = 49/25 (Indiana/Caltech) participants (mean (SD) age 24.9 (6.5) / 34.2 (4.8), from an original sample of *N* = 63 /29, prior to data-quality-related exclusions reported below). The current dataset includes predominantly males (40 Indiana, 19 Caltech) because its primary purpose is to serve as a matched control for the (mostly male) ASD participants whose data will be reported elsewhere. All subjects provided written informed consent; all experimental procedures were approved by the Institutional Review Boards of Indiana University (IU IRB) and the California Institute of Technology.

#### Unmatched (pilot) dataset

The pilot dataset was collected between 2015-2016 at Indiana University, prior to a scanner upgrade from a 3T Siemens TIM Trio to a 3T Siemens Prisma.Fit system, and is described in Byrge & Kennedy (2020). This dataset also included both typically developed adults and adults with autism spectrum disorder, and is accordingly skewed male. Only data from typically developed adults (N=25, 22 male; mean (SD) age 25.11 (4.66) years) is included in this report. All subjects provided written informed consent; all experimental procedures were approved by the IU IRB.

### Design

#### Matched datasets

Participants underwent two scanning sessions separated by approximately one week. Each session consisted of interleaved rest and video scans in a fixed order. A total of ten functional scans were collected (six video scans; four ∼16-min. resting-state scans). Table 2 presents an overview of the video scans included for each dataset. For this report, we focus most analyses on Video1 and Video2, because they were also used in the pilot dataset. For a few additional analyses, we also include the remaining four video scans. Resting-state scans were used as comparison scans for some analyses. These and the remaining functional scans will be reported in further detail elsewhere.

**Table 2.** Video scans and sample sizes for matched and unmatched datasets. Table presents video scans, initial sample sizes, and exclusions for matched datasets (IU, Cal) and unmatched datasets (Cal, Pilot). IU, Indiana. Cal, Caltech. Video1 and Video2 are the primary scans analyzed here because those video stimuli were used in all three datasets. Scan issues include technical problems (muffled sound, projector issues, missing image data) and participant sleep. Quality assurance workflow differed for the matched and unmatched datasets and MRIQC and ISC outlier exclusions were not applicable to the pilot dataset. Columns with a white background denote scans collected during the first session; columns with a gray background denote scans collected during a second session approximately one week after the first. Video scans 1-4 were all preceded by rest scans.

|  | Video1<br>movie trailers<br>~13.5 min. |  |  | Video2<br>movie trailers<br>~13 min. |  |  | Video3<br>The Office<br>~22 min. |  | Video4<br>The Office<br>~22 min. |  | Video5<br>Partly cloudy<br>~5.6 min |  | Video6<br>Bang<br>~8 min. |  |
| --- | --- | --- | --- | --- | --- | --- | --- | --- | --- | --- | --- | --- | --- | --- |
|  | IU | Cal | Pilot | IU | Cal | Pilot | IU | Cal | IU | Cal | IU | Cal | IU | Cal |
| Initial sample | 61 | 29 | 29 | 56 | 28 | 29 | 61 | 28 | 56 | 28 | 56 | 28 | 54 | 28 |
| Scan issues | 1 | 1 | 0 | 2 | 2 | 0 | 1 | 2 | 1 | 4 | 0 | 1 | 0 | 2 |
| MRIQC outlier | 5 | 1 | n/a | 1 | 1 | n/a | 4 | 1 | 1 | 0 | 4 | 0 | 1 | 1 |
| Motion | 3 | 1 | 4 | 3 | 0 | 4 | 2 | 2 | 2 | 1 | 2 | 1 | 1 | 3 |
| Registration | 1 | 0 | 0 | 4 | 1 | 0 | 2 | 0 | 3 | 0 | 2 | 1 | 3 | 0 |
| ISC outlier | 3 | 1 | n/a | 1 | 1 | n/a | 3 | 1 | 0 | 0 | 2 | 0 | 2 | 0 |
| **Final sample** | 48 | 25 | 25 | 45 | 23 | 25 | 49 | 22 | 49 | 23 | 46 | 25 | 47 | 22 |

Stimulus construction for the primary video scans (Video1 and Video2) is described in Byrge & Kennedy (2020); briefly, both scans consisted of sequences of 4-6 movie trailers collected from Vimeo (https://vimeo.com) across different genres (e.g. documentary, drama, adventure). Video3 and Video4 were different episodes of the TV sitcom “The Office (Season 1 Episode 6, “Hot Girl”; see also Byrge, Dubois, et al., 2015, and Pantelis et al., 2015; and Season 1 Episode 5, “Basketball”). Video5 was a short animated movie, Pixar’s “Partly Cloudy,” (Reher & Sohn, 2009; see also Richardson & Saxe, 2019). Video6 was an edited excerpt from the episode “Bang! You’re dead” from the television series Alfred Hitchcock Presents (1961; see also Hasson et al., 2004). Sample sizes for each video scan are reported in Table 2.

Video was back-projected onto a screen that was visible to subjects via a mirror attached to the head coil, with audio provided using Sensimetrics MR-compatible headphones. No video stimulus was provided during resting state scans (the projector was set to a black screen), and wakefulness was monitored via an MR-compatible remote eye tracker camera (Eyelink 1000+, SR Research Ltd. Ottawa, Canada). Subjects were instructed to move as little as possible and to remain awake with eyes open. Scans where problems occurred during acquisition (technical problems, such as muffled audio or issues with projector screen, or participants falling asleep) were also excluded (see Table 2).

Anatomical images were acquired following functional runs, during which participants chose to rest or watch a different video.

#### Unmatched dataset

The experimental design for this dataset is described in detail in Byrge & Kennedy (2020). Briefly, this study was also collected across two scan sessions separated by approximately one week, with interleaved rest and video scans. Only the two video scans that used the same stimuli as the primary datasets (Video1 and Video2) were included in this report. Anatomical images were collected following functional scans. See Table 2 for sample sizes.

### Data acquisition, preprocessing, and quality assessment

#### Matched dataset

MRI images were acquired using identical Siemens 3T Magnetom Prisma.Fit scanners (Siemens Medical Solutions, Natick, MA) at each site, with 64-channel head receive arrays. Scan protocols were matched across sites. Scanner software versions used were VE11B (IU) and VE11C (Caltech, and last 5 scans at IU). During functional scans, T2*-weighted multiband echo planar imaging (EPI) data were acquired using the following parameters: TR/TE 720/30 ms; flip angle = 50 °; 2.5mm isotropic voxels; 60 slices acquired in interleaved order covering the entire brain; multi-band acceleration factor of 6 (Multiband EPI sequence version R16, CMRR, University of Minnesota). Scan lengths were as follows: video 1, 1130 volumes; video 2, 1080 volumes; rest, 1355 volumes. Prior to the first functional scan, spin-echo EPI images were acquired in opposite phase-encoding directions (3 images each with P-A and A-P phase encoding) with identical geometry to the EPI data (TR/TE = 4390 / 37.2 ms; flip angle = 90°) to be used as a fieldmap to correct EPI distortions. High-resolution images of the whole brain were acquired as anatomical references (multi-echo MPRAGE, 0.9mm isotropic voxel size; TR = 2550.0 ms / TEs = 1.63 ms, 3.45 ms, 5.27 ms, 7.09 ms / TI = 1150 ms).

An upgrade to the trigger box occurred in the final months of data collection at the IU site, and this sporadically resulted in an intermittent missed trigger and delayed movie start for 35 scans. These scans were identified empirically and adjusted accordingly (see Supplemental Methods); these realignments did not influence the pattern of results reported here, which were effectively identical when conducted with the original (non-realigned) scans.

DICOM images were converted to BIDS format (Gorgolewski et al., 2016) before being run through MRIQC (v0.15.2; Esteban et al., 2017) for initial quality assessment using the functional image quality metrics (IQMs) FWHM avg, SNR, TSNR, DVARS std, and GSR. Outliers on these IQMs (the median for that data site plus or minus 1.5 times the interquartile range (IQR) for that IQM for that data site, as appropriate for the measure in question) were flagged for manual review by two of the authors (LB & DK). Following review, the consensus decision was to exclude all such flagged scans from further analyses (see Table 2).

After initial quality assessment, preprocessing was conducted using fMRIPrep (Esteban, Markiewicz, et al., 2018). The boilerplate text generated by fMRIPrep, with complete preprocessing details, is included in Supplemental Methods. Briefly, using components from ANTs (Avants et al., 2008) FSL (v. 5.0.9; FMRIB’s Software Library, www.fmrib.ox.ac.uk/fsl) and Freesurfer (v.6.0.1, Dale, Fischl, and Sereno 1999), anatomical images were bias-corrected, skull-stripped, segmented, and nonlinearly registered to MNI space. Functional scans underwent rigid-body motion correction, fieldmap-based distortion correction, and coregistration to the anatomical reference scan, and confound regressors (head motion parameters, CSF, WM, and whole-brain global signal) were computed.

For summarizing motion across a scan as well as identifying epochs of excessive motion, we computed filtered framewise displacement traces (FD_filt4_) from the fMRIPrep-computed head motion parameters, as the sums of the backwards difference across 4 TRs of motion parameters that had been filtered to exclude respiratory frequencies, as introduced by Power and colleagues (2019) and used previously for the pilot acquisition (Byrge & Kennedy, 2020). FD_filt4_ separates head motions from respiratory fluctuations in multiband acquisitions more effectively than the conventional framewise displacement computations (Power et al., 2019).

We excluded all scans with excessive motion, as identified by mean FD_filt4_ exceeding the median plus 1.5 times the IQR of the mean FD_filt4_ across all scans (including scans from ASD participants not included in the current analyses), computed separately at each site, resulting in the following exclusion thresholds: mean FD_filt4_ > 0.4808 for Indiana, mean FD_filt4_ > 0.5625 for Caltech (see Table 2). To ensure highest data quality, we also censored time points surrounding excessive motions: 10 frames before and 30 frames after any frame with FD_filt4_ > 3.75mm; censored time points were treated as missing data in all analyses, and the inclusion or exclusion of censored data did not influence the overall pattern of results.

All reports generated by fMRIPrep were inspected by two independent reviewers (two research assistants, one based at each site, trained to conservatively flag any potential issues with anatomical and functional scans and their alignment). All reports flagged by both research assistants were then independently reviewed by both LB & DK and a consensus decision was reached about whether to include or exclude all such flagged scans from the current dataset (see Table 2).

Subsequent preprocessing used *xcepengine* version 1.2.1; detailed in detail by Ciric and colleagues (2018). We used the “fc-24p_gsr” pipeline optimized for functional connectivity processing; this configuration is publicly available at https://github.com/PennBBL/xcpEngine/blob/master/designs/fc-24p_gsr.dsn. Briefly, functional data was demeaned and detrended, aligned to the anatomical reference scan, and bandpass-filtered within the range 0.08-0.001 Hz using a Butterworth filter. Then, 36 confound regressors (6 head motion parameters, CSF, WM, global signal, their backwards differences, and then the squares of those 18 traces; all temporally filtered in the same way as the data) were regressed from the data, and then the residuals were spatially smoothed with a 2.54mm filter, and then used as the “cleaned” data.

For the primary datasets, we examined average BOLD timeseries across several different atlases. We focused exclusively on cerebral cortex, excluding the cerebellum and all subcortical structures, following the largely cortico-centric focus of the inter-subject synchrony literature. The primary atlases used were different parcellation scales of the Schaefer atlas (Schaefer et al., 2018), which subdivides the intrinsic functional connectivity-based Yeo network parcellation of the cortex (Yeo et al., 2011) into 100, 200, 400, 600, 800, and 1000 cortical regions. We also examined the structural Harvard-Oxford Atlas distributed with FSL, which had been previously used to parcellate the pilot dataset. We restricted our analysis of the Harvard-Oxford parcellation to cortical regions of interest (ROIs; 96) only, for consistency with the Schaefer cortical parcellation. For all atlases, we obtained ROI timeseries for each region as the mean of the “cleaned” BOLD signal across all voxels in the given region, at each time point.

As an additional data quality assessment, for all video scans, we examined BOLD time series from primary visual cortex and primary auditory cortex in each hemisphere (using the Harvard-Oxford parcellation), in order to identify and exclude scans where technical problems with the stimulus presentation or visual or auditory aspects of the stimulus occurred but were not noted at the time of scanning (e.g., headphone or projector failure, or misalignment between the start of image acquisition and the video). We approached this cautiously and conservatively, because similarity of BOLD time series among scans is also our measure of interest for this report; but, at the same time, extremely low similarity to other participant timeseries in primary sensory areas during long video scans is an indicator that something has gone wrong in the scan acquisition process. Therefore, separately within each dataset, for each video scan, and for each of the four primary sensory regions of interest, we computed pairwise correlations among all participant time series, and computed the median minus three times the interquartile range of median pairwise correlations for each participant as a threshold to identify extreme outlier values suggestive of equipment issues. We excluded scans for which the median pairwise correlation was below this data-driven threshold in at least one of the sensory regions (see Table 2).

#### Exploratory pilot dataset

This acquisition and preprocessing pipeline is described more completely in Byrge & Kennedy (2020). Briefly, images were collected using a 3 T Magnetom Tim Trio system (Siemens Medical Solutions, Natick, MA) with 32-channel head receive array, running software version VB17. T2*-weighted multiband EPI data was acquired using the following parameters: TR/TE = 813/28 ms; 1200 volumes; flip angle = 60°; 3.4 mm isotropic voxels; 42 slices acquired with interleaved order covering the whole brain; multi-band acceleration factor of 3. Gradient-echo EPI images (10 images each with P-A and A-P phase encoding; TR/TE = 1175/39.2 ms, flip angle = 60°) were used as fieldmaps for EPI distortion correction. High-resolution T1-weighted images of the whole brain (MPRAGE, .7 mm isotropic voxel size; TR/TE/TI = 2499/2.3/1000 ms) were acquired as anatomical references.

Data were preprocessed using an in-house pipeline using FSL (v. 5.0.8; FMRIB’s Software Library, http://www.fmrib.ox.ac.uk/fsl), ANTs (v2.1.0; Avants et al., 2011), and Matlab_R2014b (www.mathworks.com, Natick, MA, USA). Preprocessing steps included rigid-body motion correction, fieldmap-based geometric distortion correction, non-brain removal, weak highpass temporal filtering (>2000s FWHM) to remove slow drifts. Denoising was preformed using FSL-FIX (Salimi-Khorshidi et al., 2014) followed by mean cortical signal regression in a second step (effectively the same as global signal regression, but using the signal across the cortex rather than whole brain; Burgess et al., 2016), with the residuals analyzed as the “cleaned” data. Volumetric registrations were conducted using FSL and ANTs, using a combined affine and diffeomorphic transformation matrix. Region of interest (ROI) timeseries using the Harvard-Oxford Atlas distributed with FSL were obtained as the weighted mean signal from the “cleaned” BOLD signal across voxels within each of the 110 ROIs.

As these data were collected using a different repetition time (TR) than the primary dataset (813 ms vs. 720 ms), the final preprocessing step for this report was to resample these time series to match the faster sampling rate of the primary dataset. There are many possible ways to perform such resampling; here, we used Fourier method resampling as implemented in scipy.signal.resample.

Differences between the primary and exploratory pilot unmatched acquisitions are summarized in Table 1.

### Data Analysis

Naturalistic fMRI data analysis requires evaluating the similarity of brain response time series. Here, we examined similarity of brain responses across sites at two distinct levels: similarity of group-average time series from each site (Fig. 1A), and similarity of pairs of individual subject time series across sites and within each site (Fig. 1B; pairwise inter-subject correlations; ISC; Hasson et al., 2004). At the group level, within each site, we used median time series across subjects within each brain region of interest to isolate the common brain response pattern while reducing the influence of various forms of noise. We take these measurements of across-site similarity at both levels as our measures of cross-site consistency.

**Figure 1.**
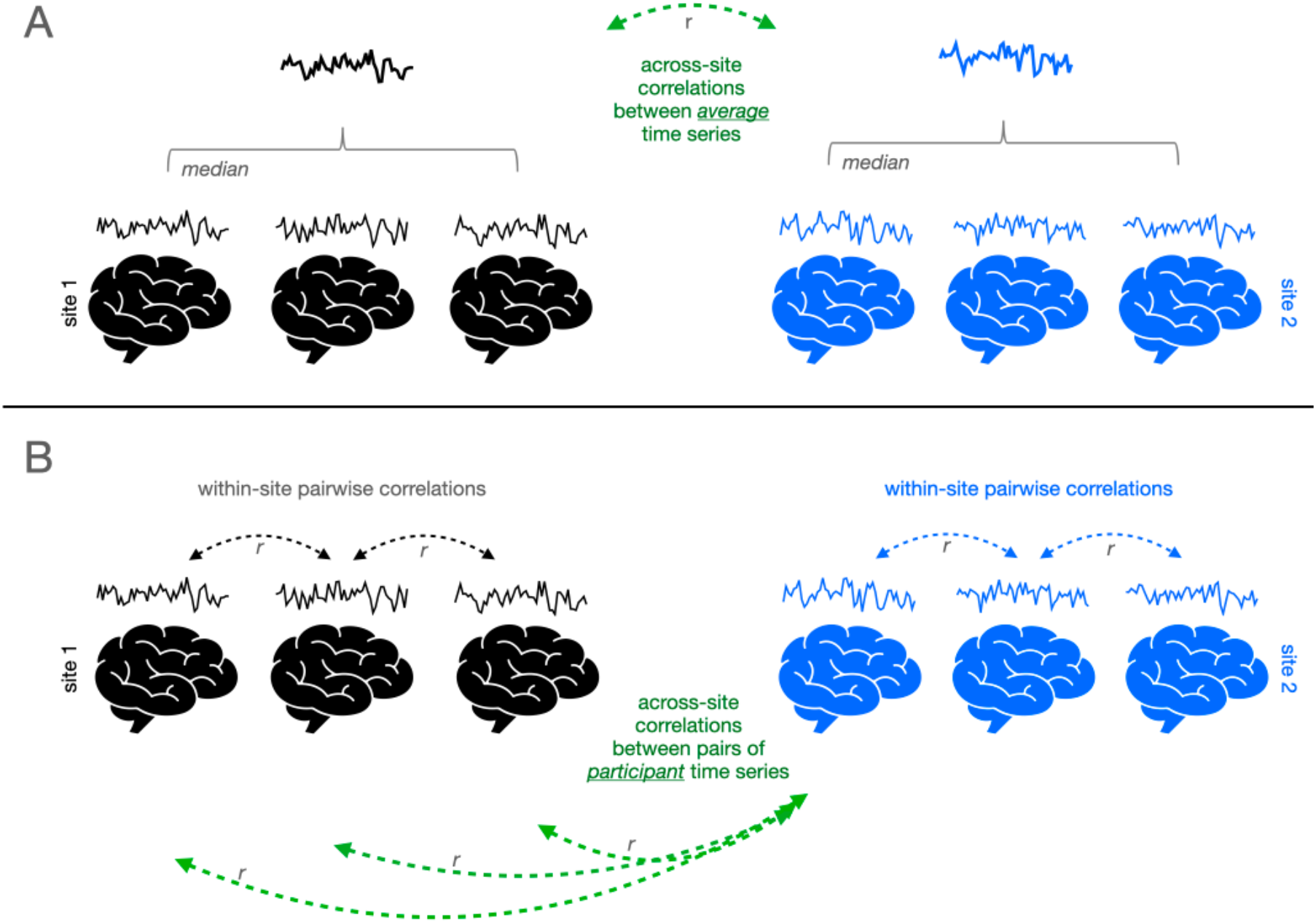
Schematic of approach for examining consistency of video-evoked brain responses across sites (black and blue) at the level of the group (A) and individuals (B). Example individual time series depict the fMRI BOLD signal averaged across a given region of interest, across the duration of the video. To examine consistency across sites at the group level (A), the average of all these individual time series is computed for each site (bolded timeseries), and then the correlation between those average site-level time series is computed (green arrow). To examine consistency across sites at the individual level (B), correlations between pairs of time series from individual participants at different sites are computed (green arrows), and for some analyses compared to correlations between pairs of participant time series from the same site (black arrows, or blue arrows).

Unless noted, analyses are repeated on the two primary video trailer scans (Video 1 and Video 2) that were used as stimuli in each of the datasets. Most analyses are conducted across multiple spatial scales using different granularities of the Schaefer parcellation (Schaefer et al., 2018).

#### Statistical comparisons

##### Consistency of brain responses across sites

As a first broad characterization of the extent to which there is shared signal across sites, we measured similarity between median time series for each site using Pearson correlations. We considered a brain region (ROI) to be responding more consistently across sites during the same video than expected by chance if the correlation was statistically significant following FDR-correction for the granularity of the parcellation in question. (Note that the patterns of results were effectively the same when using a non-parametric, bootstrapped procedure).

For individual-level analyses, for each scan and in each brain region, we computed pairwise inter-subject correlations (ISC) as the Pearson correlation among pairs of individual brain responses for all pairs of subjects. We also computed pairwise similarity across different scans (e.g., between Video1 and Rest) for use in null models as described below. We used non-parametric statistical comparisons for this level of analysis, following the recommendations of Chen and colleagues (2016) for pairwise ISC, and pooled pairwise correlations within and/or across sites without collapsing at the individual level.

To evaluate whether a brain region responded more consistently across sites than expected by chance at the individual level, we examined only inter-subject correlations between pairs of participants from different sites, and asked whether the magnitudes of those correlations were greater when both participants were watching the same video than when one participant was watching the video and the other participant was resting. We compared correlation magnitudes (absolute values) to avoid exaggerated influence from very small negative correlations, which were expected between video and rest scans. We held the comparison scans fixed at each site, yielding two observed median correlation differences. Specifically, within each brain region, for video scan V and rest scan R, across all across-site pairs of Indiana participants I*i* and Caltech participants C*j*, we obtained: Δr_Indiana_= *median*(| *r*(V_I1_,V_C1_) |, …, | *r*(V_Ii_,V_Cj_) |) – *median*(| *r*(V_I1_, R_C1_) |, …, | *r*(V_Ii_, R_Cj_) |) and Δr_Caltech_ = *median*(| *r*(V_I1_,V_C1_) |, …, | *r*(V_Ii_,V_Cj_) |) – *median*(| *r*(R_I1_, V_C1_) |, …, | *r*(R_Ii_, V_Cj_) |). We then permuted scan type labels (Video vs Rest) 10000 times and computed this same measure to establish a null distribution of median differences, and obtained a one-sided empirical *p*-value (corrected to avoid bias due to finite sampling, Davison & Hinkley, 1997) of observing Δr_Indiana_ and Δr_Caltech_ by chance. To address multiple comparisons, we applied FDR correction within each parcellation. We conservatively considered a region as responding more consistently than expected by chance only if it survived FDR correction for both Δr_Indiana_ and Δr_Caltech_ (note also that results were effectively the same for both cases).

##### Differences between sites

After examining consistency of brain responses between sites, we examined differences. To do this, and to get estimates of variance, we continued at the individual subject level, because group average time series mitigate or even eliminate noise that might be unique to a given site or scanner. Differences between datasets would manifest as differences in within-site similarity vs. across-site similarity. Therefore, for each video, site, and brain region, we computed the observed difference in within vs. across-site ISC as the difference between the median ISC among all subjects at the site in question and the median ISC among all different-site subject pairs. As before, we computed this measure separately for each site, because levels of within-site similarity could be different. We then computed the empirical p-value of observing this median difference as before using a permuted null distribution constructed by shuffling site labels 10000 times and computing the permuted median difference. For this analysis, we conservatively used α = 0.05 with no correction for multiple comparisons, in order to increase our sensitivity to detect potential differences between datasets (at the expense of likely false positives). Finally, to contextualize the magnitudes of differences between datasets, we also compared distributions of within- and across-site ISC values using Mann-Whitney rank sum tests and report the common-language effect sizes (CLES; Vargha & Delaney, 2000) across ROIs. Common-language effect sizes in the *within – across* direction reported here reflect the proportion of pairs of observations in which within-site pairwise ISC is higher than across-site pairwise ISC. CLES of 0.5 indicates no effect, and CLES of 0.56, 0.64, and 0.71 roughly correspond with Cohen’s *d* values of 0.2, 0.5, and 0.8, indicating small, medium, and large effect sizes (Ruscio, 2008). (Note that CLES below 0.5 would indicate higher across-site ISC than within-site ISC with comparable interpretations, e.g. CLES of 0.44 would reflect a small effect.)

### Results

#### 1. SIMILARITY

##### 1.1. Consistent group-level brain responses are evoked by the same videos at different sites

Average brain response time series across a group of participants should capture common patterns of stimulus-evoked brain function while mitigating the effects of physiological noise, scanner noise, and individual differences in brain functioning. The hope, for generalizability of vfMRI studies, would be that these group-level brain responses would be largely similar across sites (as depicted in Fig. 1A), especially when acquisitions and processing are matched as closely as they are in these primary datasets. Figure 2 shows correlations between the median time series across IU participants and the median time series across Caltech participants in each brain region while watching the videos, under different spatial scales of the Schaefer atlas (Schaefer et al., 2018), which subdivides the intrinsic connectivity-based Yeo network parcellation of the cortex into 100, 200, 400, 600, 800, and 1000 regions. As is evident, average brain responses across Indiana participants were highly similar to average brain responses across Caltech participants, during both video scans, across all parcellation scales examined. Highly similar brain responses were not limited to the primary sensory areas expected to be driven by the stimulus (c.f. visual and auditory timeseries shown in Fig. 2) but extended throughout the cortex (c.f. association timeseries shown in Fig. 2).

**Figure 2.**
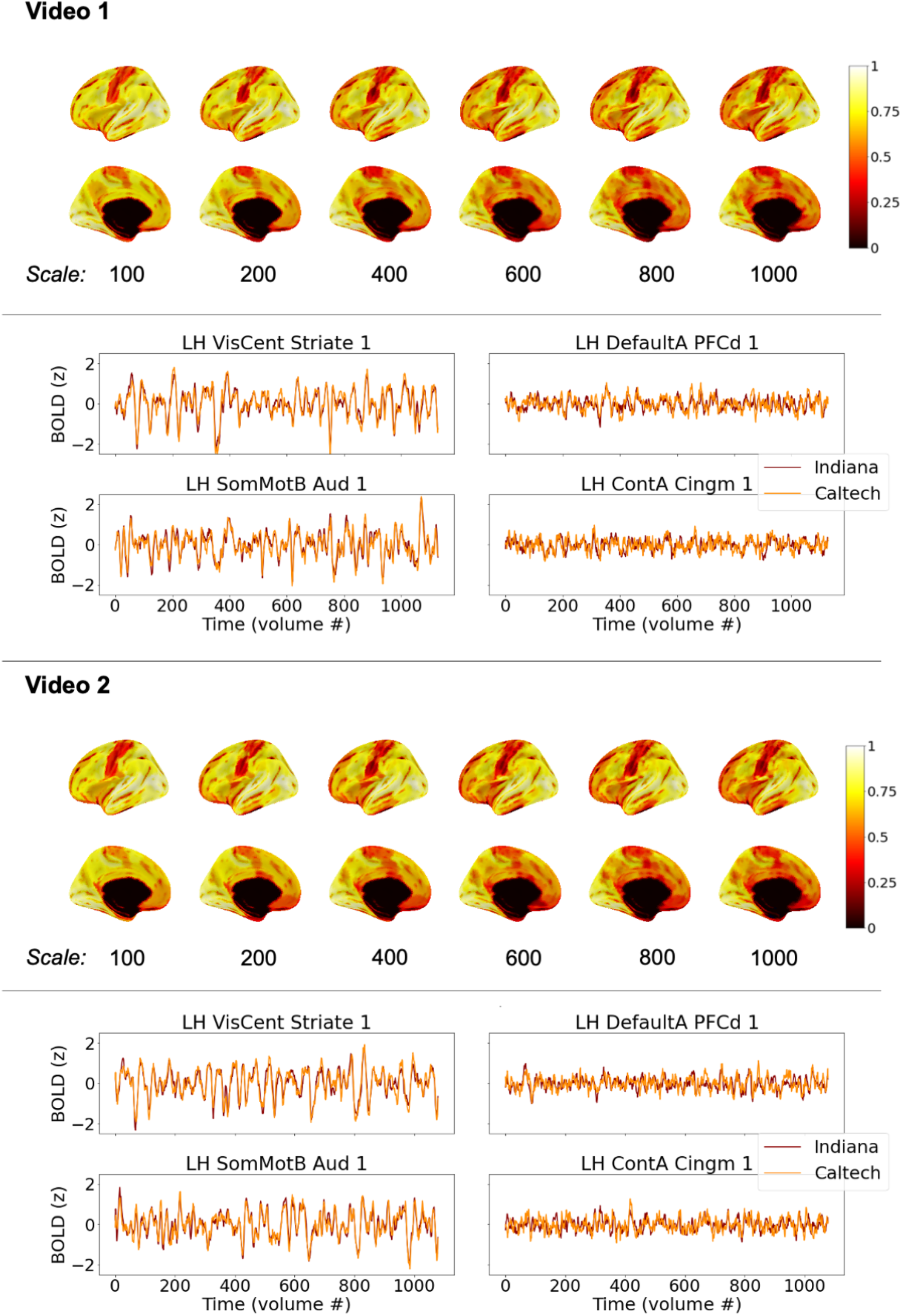
Consistency of average group-level brain responses across sites while participants watched the same videos (see Fig. 1A), for matched datasets. Brain visualizations depict correlations between median time series across all participants at each site, in each brain region, under different scales of the Schaefer parcellation (100-1000 ROIs). The Schaefer parcellation is a cortical parcellation; black along the midline in medial views here and elsewhere indicate missing data, not low correlations. Line plots depict median timeseries across participants at each site in primary sensory areas (left) and association areas (right) using the 400-region Schaefer parcellation during Video 1 (top) and Video 2 (bottom). All figures depict the left hemisphere; the pattern of results for the right hemisphere is effectively the same.

##### 1.2. Consistent group-level brain responses are found through most of the cortex

As is evident in Figure 2, highly similar group-level brain responses across scanners were not limited to the coarser parcellations. As parcellation granularity increases, though, some correlation magnitudes decrease – as expected, as regional timeseries approach voxel-level timeseries with correspondingly reduced spatial smoothing – to the extent that it becomes unclear by eye whether brain response similarity across scanners exceeds chance levels in some brain regions. In all parcellations, the median time series at each site was more correlated than expected by chance in more than 99% of brain regions. However, the size of these effects varied across the brain, as can be seen on the color axis, and in some association areas, significant correlations between group-level brain responses at each site were quite small (around *r* = 0.1, after FDR correction). One might expect such a weak shared signal to be easily dominated by other factors (e.g. endogenous processing, scanner noise, registration inaccuracies) if not for averaging across multiple scans (i.e., participants).

Brain regions that did not respond consistently across sites at the group level were detected only in the finer parcellations and included parts of the temporal pole (an area prone to susceptibility artifacts) as well as part of medial prefrontal cortex during one scan.

##### 1.3. Participant-level brain responses are consistent across sites throughout most of the cortex

Average time series across groups of subjects are effective at isolating common response patterns by dampening down individual variability, but they are not representative of any individual brain’s functioning and could minimize potential differences in noise properties across scanners. Thus, next we examined consistency of individual brain responses among pairs of participants as they watched the same videos (as depicted in Fig. 1B). Figure 3 (see also Supplemental Figure 1) shows the median of these pairwise inter-subject correlations across sites in the center column, along with pairwise ISCs within each site (left and right columns) for comparison. As expected, based on increased noise and individual variability in participant time series, the range of correlations is shifted lower than in Figure 2, but the general conclusion remains the same: consistent brain responses across sites are widespread throughout the majority of the cortex. Nearly all ROIs (>90% in all six cortical parcellations; ranging from 98% of ROIs in the 100-ROI parcellation and 91% in the 1000-ROI parcellation) were more similar between cross-site pairs of subjects watching the same video than expected by chance, using a null distribution comprised of pairwise brain response similarity in which one participant watches this same video and the other undergoes a resting state scan.

**Figure 3.**
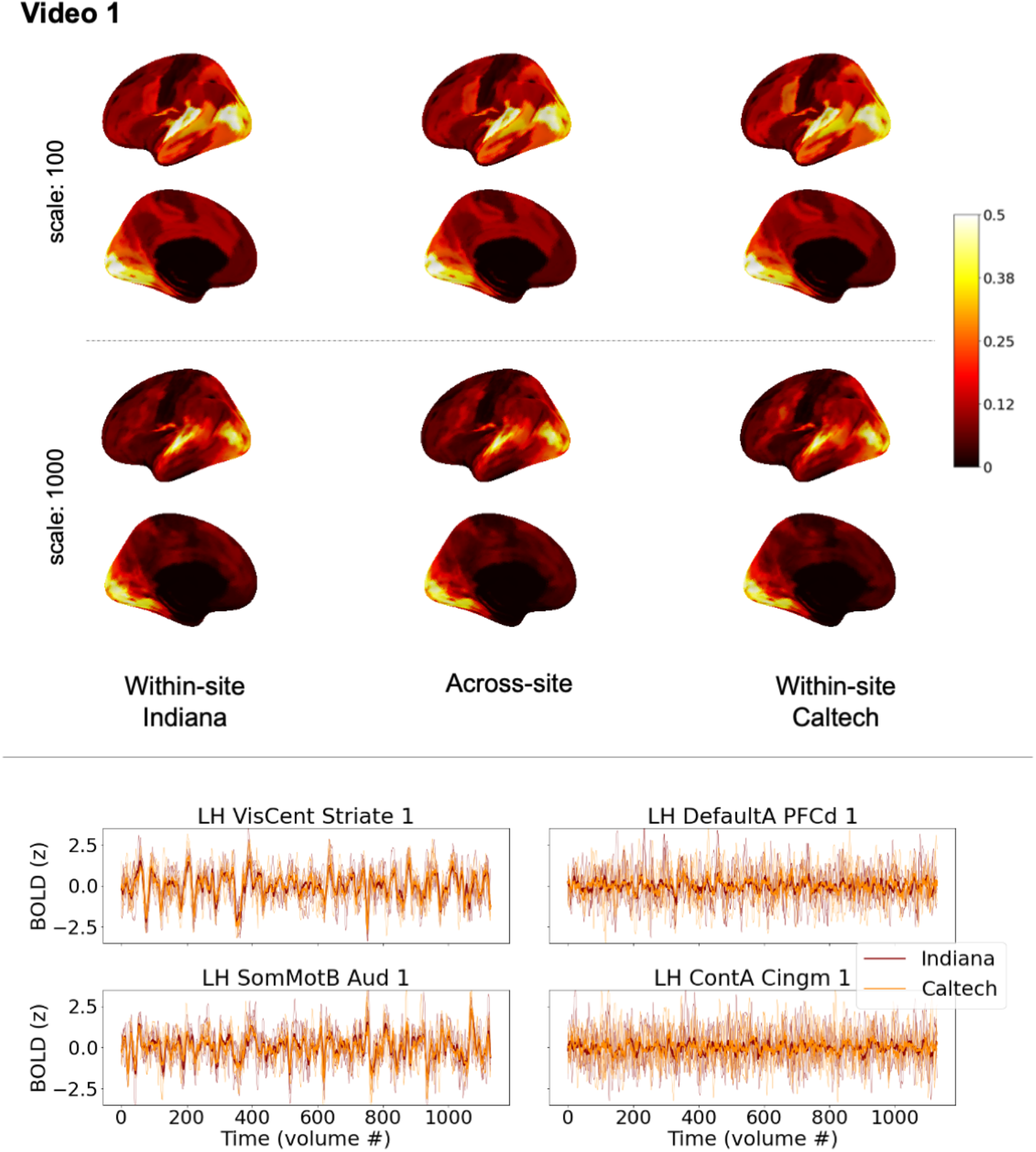
Similarity of individual participant brain responses within and across sites during Video1 (see Fig. 1B) for matched datasets. Top: brain maps depict magnitudes of medians of pairwise correlations between participant brain response time series in each region, for the most coarse (top) and the most fine (bottom) scales of the Schaefer cortical parcellation. The left and right columns show correlations among pairs of participants at the same site (left: Indiana; right: Caltech). The center column shows correlations among pairs of participants spanning different sites. While absolute values are depicted here for readability, nearly all median correlations were positive, except one temporal pole ROI with a near-zero median correlation of -0.0036. As in Figure 2, black along the midline in medial views indicates missing data, not low correlations. See also Figure 4 (bottom panel, top plot) for a line version of this same data, and Supplemental Table 2 for characterization of differences and effect sizes. Bottom: line plots depict timeseries from 5 randomly-selected participants at each site in primary sensory areas (left) and association areas (right), along with the site-level median time series shown in Figure 2. These line plots use the same mid-scale parcellation as Figure 2 (Schaefer 400x17). See also Supplemental Figure 1 for the equivalent figure for Video2, which is similar but supplemental for space purposes.

It is important to note, though, that above-chance similarity relative to resting state does not imply a large effect size, and the median across-site pairwise ISCs in some ROIs that responded at above-chance levels could be exceedingly small, even below 0.01. In other words, although the common stimulus explained some proportion of variance in these time series (and a vanishingly small proportion for these smallest correlations), most variability is left unexplained – and thus left to be explained in future studies of stimulus-level, contextual, state-level, physiological, and phenotypic factors underlying these individual brain responses (see e.g., Chang et al., 2021). This is the case even for the primary sensory areas most strongly driven by the stimulus, where median across-site pairwise correlations could exceed 0.5, which still leaves around 75% of the variance unexplained by the common stimulus. Inspection of the randomly selected example time series in Figure 3 (bottom) and Supplemental Figure 1 suggests the possibility that some specific moments of the stimulus may drive relatively instantaneous similarity amid otherwise dissimilar brain responses in some of the association areas that have low ISC that nonetheless exceeds chance (c.f. Figure 3, PFCd), but future work will be needed to examine that possibility directly.

##### 1.4. Consistent group- and individual-level brain responses are evoked by a variety of video stimuli

Participants in the primary dataset watched different video stimuli sampling a variety of genres: movie trailers (Videos 1 and 2; the main stimulus throughout this manuscript), complete episodes of TV sitcoms (“The Office”; Videos 3 and 4), an animated short film (“Partly Cloudy”; Video 5), and a black-and-white Alfred Hitchcock film (“Bang! You’re dead”; Video 6). As is already apparent in Figure 2, cross-site consistency at the group level was high for both Video1 and Video2 despite varying stimulus content – each of those videos consisted of sequences of trailers for entirely different movies. Figure 4 (top) shows the correlations between group-average timeseries at both sites for all six video scans in the primary dataset, using the coarsest and finest parcellation scales. As is evident, high cross-site similarity between average time series is a feature of all the video stimuli used, and not limited to movie trailers alone. Note that while the maps look similar – qualitatively, regions that responded highly consistently across scanners during one video also responded consistently during the other video – the patterns are also not identical across different videos. These differences could reflect differences in stimulus content. For instance, reductions in cross-site similarity can be observed in some temporal and frontal regions during the largely silent animated film (Video5). Notably, episodes of The Office were chosen so as to emphasize social features in the video, and cross-site consistency in medial prefrontal cortex appears elevated in Videos 3 and 4 relative to the other scans, potentially reflecting the increased social processing demands of the stimulus. While these video scans also differed in length, scan length did not appear to be the main driver of these differences (see also Supplemental Figure 2, which presents an alternate version of this figure that was randomly downsampled to address length differences, but shows largely similar patterns).

**Figure 4.**
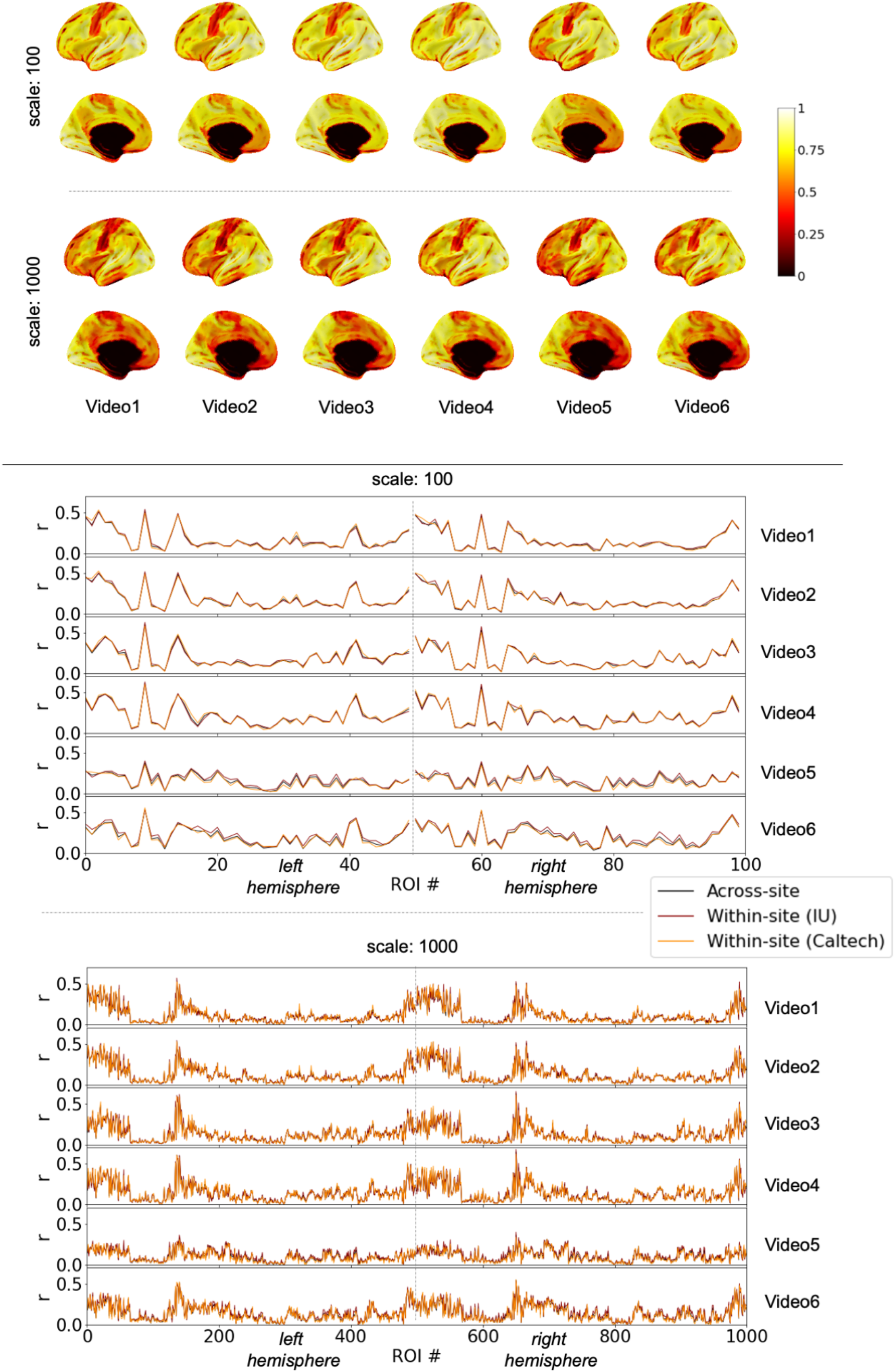
Consistency in brain responses across sites while participants watched the same video, across a variety of different videos, in the matched datasets. Top: brain maps show cross-site consistency at the group level (as in Figure 2, see also Fig. 1A). Only parcellation scales of 100 and 1000 ROIs are shown for space considerations. As in Figure 2, black along the midline in medial views indicates missing data, not low correlations. Scan lengths vary across different videos. See also Supplemental Figure 2 for a version of this figure that randomly downsamples to equate for scan length. Bottom: line plots show median pairwise ISC among pairs of participants within each sites and across sites (see Fig. 1B), for the most coarse and most fine parcellation scales. Please see Supplemental Table 1 for ROI labels, which are omitted for readability. The line plots for Video 1 and 2 are the same values plotted on brains in Figure 3 (top) and Supplemental Figure 1. Note that individual data points are connected with a line to facilitate comparing overall patterns, but these plots are not time series. Rather, each data point reflects median similarity across pairs of timeseries. When values for a given ROI are similar across different scans (e.g. x = 60 for top two line plots), that reflects comparable levels of similarity across entirely different brain response time series for different videos.

Figure 3 (top) and Supplemental Figure 1 showed high levels of cross-site consistency at the level of individual subject pairs for Videos 1 and 2. Figure 4 (bottom) shows median across-site (in black) and within-site (in color) pairwise inter-subject correlations for each of the video scans in each ROI. One of the colored lines corresponds to the values that would be projected onto a brain map if the data had been collected at a single site (e.g., the red line for Video1 in Figure 4 (bottom) is the same as the Indiana within-site ISC map in Figure 3, upper left). While the colored lines show small intermittent deviations above and below the black line, the larger take-away is that the lines track one another closely. The pattern of median within-site ISC across ROIs for one site is highly correlated with the pattern of within-site ISC for the other site, and both quantities are highly correlated with median across-site ISC (all *r* > 0.91 across all videos and all parcellations). In other words, brain responses for pairs of subjects at different sites are about as similar as pairs of subjects at the same site. This close tracking is apparent both for brain regions that are more evoked and less evoked by the stimuli, as well as for different video stimuli that drive higher and lower ISC values in the same brain regions (for instance, c.f. *x* = 41 (part of left hemisphere temporal lobe) for Video5 vs. the other video scans, for the center panel with the 100-scale parcellation). Cross-site consistency of brain responses for this set of stimuli is neither limited to a few sensory regions that are most strongly driven by the video, nor a subset of video stimuli that drive the brain especially strongly, but is instead apparent throughout the different stimuli used here.

### 2. DIFFERENCES

#### 2.1. Differences across sites are minimal when acquisitions and processing are matched

After establishing that brain responses across the cortex are indeed consistent across sites, the natural question is to ask about differences. If there were no differences between datasets, an individual scan would be just as similar to other scans at the same acquisition site as it is to other scans from a different acquisition site. Arguably, site-level differences could manifest as either increased or decreased similarity among participants at the same site, depending on noise properties. We thus evaluated potential site differences by testing whether within-site pairwise similarity for either site differed from across-site pairwise similarity in any brain region, by comparing the observed differences of the medians to a permuted null distribution in which site labels were shuffled. Because levels of within-site consistency need not be the same for both sites, and therefore differences between within-site and across-site similarity could differ, we considered a brain region to have a site difference if such a difference was observed for either site, and not necessarily for both sites. For this analysis, to conservatively increase sensitivity for detecting any potential differences, we did not correct for multiple comparisons, and thus some false positives are likely.

For all video scans, and for all parcellations, the majority of brain regions had no site differences at this conservative threshold. Proportions of brain regions that did have site differences ranged (across parcellation scales) as follows: Video1, 0.13-0.17; Video2, 0.09-0.15; Video3, 0.06-0.11; Video4, 0.06-0.13; Video5, 0.23-0.32; Video6, 0.35-0.47. As noted above, these proportions are likely to be an overestimate. Differences can be observed in Figure 4 (bottom) as gaps between the black and colored lines. Differences are generally small relative to the level of ISC, and are found in regions including those that are strongly driven by the stimulus (e.g., *x* = 53, part of the right hemisphere peripheral visual network, for the center panel with the 100-scale parcellation). Supplemental Table 2 summarizes differences in the distributions of within-site and across-site ISC values. For all videos and all parcellations, median differences between within- and across-site ISC values across ROIs were small (<0.03), with median common-language effect sizes (CLES) across ROIs corresponding with small effect sizes. Maximum CLES across ROIs reflected small or medium effects depending on the video and parcellation (maximum CLES from 0.57 to 0.67), but never large effects. Such differences could arise due to different levels of individual variability or effects of scanning equipment per se (or both). Regardless of the sources, it is important to note that when datasets are closely matched, as they are in this primary acquisition, most cortical regions did not show site differences even at this sensitive threshold, the effect sizes of site differences were predominantly small, and, as noted earlier, patterns of within-site ISC for each site were highly correlated with one another and with the pattern of across-site ISC.

#### 2.2. When acquisitions are not matched, differences become more apparent, despite still-widespread consistency

In an exploratory comparison we also examined cross-site differences between the primary Caltech dataset and a pilot dataset (“Pilot”) also collected at Indiana University prior to a scanner upgrade. These unmatched datasets were collected using different scanner models, protocols with numerous differences, and different preprocessing approaches (Table 1), but using the same scanner manufacturer (Siemens) and field strength (3T). As these acquisitions were not designed to systematically test the effects of varying all these parameters, it will not be possible to disentangle the specific sources of any differences identified.

Nonetheless, we include this comparison as somewhat more representative of real-world differences between pre-existing datasets collected as participants watch the same video stimulus.

Figure 5 (center) depicts consistency between these two unmatched datasets at the level of median time series at each site, along with consistency between the two primary matched datasets (left), and the difference of these quantities (right), for comparison. Median time series from all three sites are presented as well. The same general pattern of results from Figure 2 is evident even though the Caltech and Pilot acquisitions are unmatched: high similarity across datasets at the group average level while participants watch the same video stimulus. Despite this high similarity, though, it is also visually apparent that similarity between the unmatched acquisitions is reduced, relative to the matched acquisitions.

**Figure 5.**
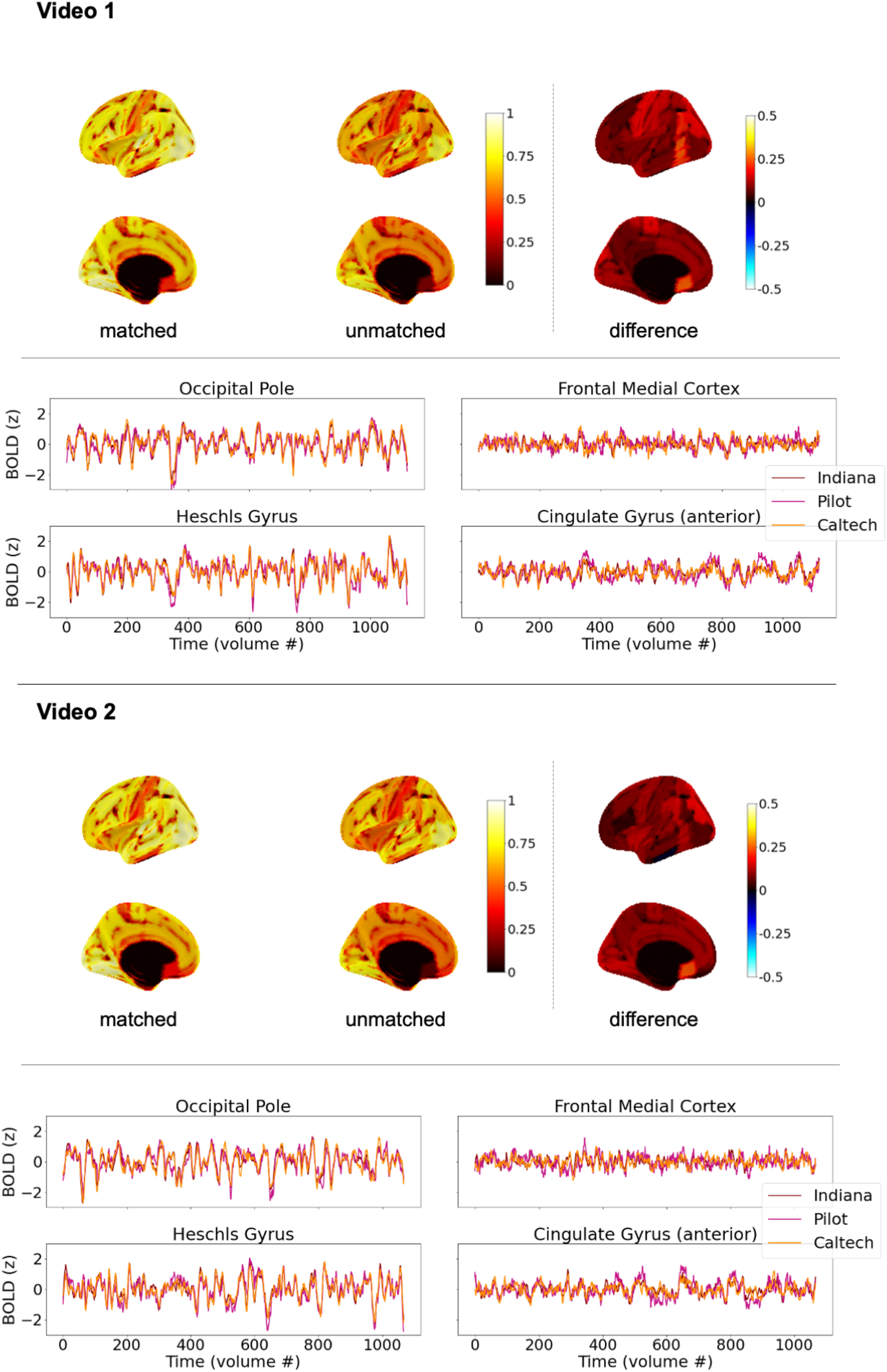
Exploratory comparison of brain response consistency for matched datasets and unmatched datasets, at the level of average time series (see also Fig. 1A), for Videos 1 and 2. Left and center brain maps show correlations between median time series for all participants at each site (as in Fig. 2, top); right shows the difference of the two maps. Matched datasets are Indiana and Caltech (as in Fig. 2); unmatched datasets are Pilot and Caltech. Black along the midline in medial views indicates missing data, not low correlations. Similarity of average time series across datasets is high in general, but highest when acquisitions are matched. Time series figures show median time series for each site in sensory areas (left) and association areas (right). Alll panels use the Harvard-Oxford 96-ROI cortical parcellation.

Because potential differences between datasets are expected to manifest most strongly within individual subject data, we tested for differences at the level of pairwise ISCs in the same way as described previously, by testing whether within-site similarity differed from across-site similarity for either site. Pairwise ISCs within and across each of these unmatched datasets are mapped in Figure 6 (top, and Supplemental Figure 3), and also presented as line plots (bottom) to facilitate comparison. Differences in ISC levels are visible, with within-site similarity for the Pilot dataset appearing elevated. To test for differences, as before, to be conservative, we did not correct for multiple corrections, and considered a brain region as having a site difference if differences were observed for either site (and not necessarily both). In contrast to the previous results for the matched datasets (IU vs. Caltech, see 2.1), here, when datasets were unmatched in numerous ways, we observed site differences in most brain regions: 89.6% of regions for Video1, and 97.9% for Video 2.

**Figure 6.**
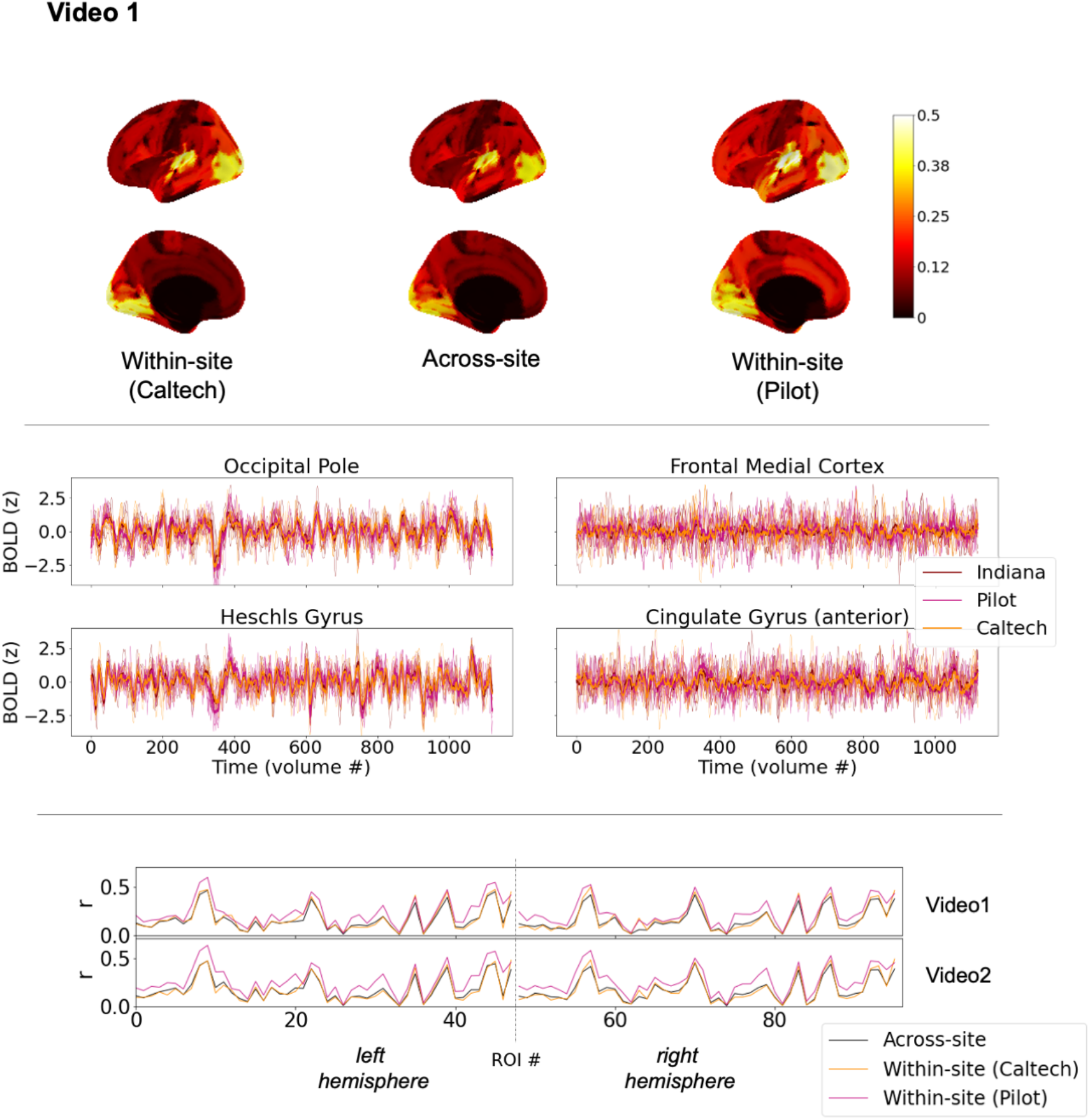
Exploratory comparison of brain response consistency across unmatched datasets, at the level of individual time series (Fig. 1B), for Video1. Top: brain maps depict magnitudes of medians of pairwise correlations between participant brain response time series in each region. The left and right columns show correlations among pairs of participants at the same site (left: Caltech; right: Pilot). The center column shows correlations among pairs of participants spanning different sites (as in Fig. 1B, green). Black along the midline in medial views indicates missing data, not low correlations. Center: Time series plots show five randomly-selected individual time series for each site in sensory areas (left) and association areas (right), along with the median time series across all participants at that site superimposed in bold. Bottom: Within- and across-site ISC values from top panel (Video1) and Supplemental Figure 3 (Video2) presented as a line plot, to facilitate comparison. See text for summary of differences and effect sizes, and see Supplemental Table 1 for ROI labels, which are omitted for readability. As in Figure 4 (bottom), individual data points are connected with a line, but these plots are not time series. Rather, each data point reflects median similarity across pairs of timeseries. When values for a given ROI are similar across the two different scans, that reflects comparable levels of similarity across entirely different brain response time series evoked by different videos. All panels use the Harvard-Oxford 96-ROI cortical parcellation.

As can be seen in Figure 6, differences between median within-site and median across-site ISC varied across ROIs and varied by site. They appear relatively minimal for the Caltech dataset but more noticeable for the Pilot dataset, and they are not homogenous across the brain. For instance, elevated within-site similarity in the Pilot dataset was found throughout the superior temporal lobe extending into the temporoparietal junction (c.f. Figure 6, *x* = 9 and 57, left and right posterior superior temporal gyrus), but much less so for many visual areas (c.f. *x* = 39 and 87, left and right occipital fusiform). For the within-Pilot vs. across-site ISC comparison (pink vs. black lines), these differences ranged across ROIs from 0.008-0.2 (median 0.06; IQR 0.06) for Video 1 and from 0.006-0.24 (median 0.08; IQR 0.5) for Video2. CLES for these differences ranged from 0.52-0.92 across ROIS (median 0.66; IQR 0.11) for Video1 and from 0.52-0.95 (median 0.71; IQR 0.11) for Video2. For the within-Caltech vs. across-site ISC comparison (orange vs. black lines), the median difference across ROIS ranged from -0.05-0.12 (median 0.02; IQR 0.035) for Video1 and -0.06-0.1 (median -0.006; IQR 0.03) for Video2. CLES ranged from 0.32-0.76 (median 0.49; IQR 0.1) for Video1 and from 0.32-0.81 (median 0.47; IQR 0.1) for Video2. In contrast to the matched datasets, then, quantitative comparisons of pairwise ISC levels within- and across-sites can reveal differences with medium-to-large effect sizes spanning ROIs.

Due to the numerous factors that vary between these unmatched datasets (Table 1), it is not possible to pinpoint the exact cause(s) of the elevated within-site similarity in the Pilot dataset; disentangling these factors is beyond the scope of the current project and a question for future targeted new acquisitions. Nonetheless we present these comparisons as a case study showing how similarity across and within sites can vary when datasets using the same video stimuli are unmatched.

And while quantitative differences in ISC levels were prevalent in comparing these unmatched datasets, it is important to observe that qualitatively, the pattern of ISC remained similar across sites and within each site. Figure 6 shows that all three lines increase and decrease in tandem, and indeed they are all highly correlated for both video scans (all *r* > 0.92, *p* < 0.0001). So, while levels of ISC can differ considerably when datasets are unmatched, ROIs with higher ISC at one site also have higher ISC at the other site and across-sites, and vice versa.

Altogether, these results indicate that differences in brain responses across sites are more readily apparent when datasets are unmatched, and can be considerable and non-homogeneous across the cortex – but, despite these quantitative differences, video stimuli drive qualitatively consistent patterns of brain responding across sites even when numerous acquisition, processing, hardware, and participant details vary freely.

## Discussion

We find that video fMRI paradigms evoke robustly similar brain responses across different sites and samples of subjects, with consistent brain responses found through most of the cortex. When datasets are matched closely, such that scanner manufacturer, model, imaging protocols, and preprocessing details are the same at each site, differences in brain responses between datasets are minimal. When datasets are unmatched, such that scanner model and acquisition and processing details vary more freely, differences are more prevalent, especially in pairwise comparisons of individual data. Nonetheless, consistency of brain responses across unmatched datasets remains high, although attenuated relative to matched datasets.

In the matched datasets, at the level of group-average time series, we find that most regions of the brain (>99%) respond similarly across sites, and this nearly cortex-wide similarity is observed across parcellation granularities (from 100 to 1000 ROIs) – it is not an artifact of using a coarse parcellation and therefore spatially smoothing across large swaths of cortex. We find comparable results at the level of individual time series similarity, albeit with the reduced correlation magnitudes expected from pairwise correlations. Procedures adjusting for individual differences in functional specialization and hemodynamic responses (Haxby et al., 2020, Dubois & Adolphs, 2016) could be employed in the future to potentially reveal even higher similarity across sites.

Across parcellations, regions with consistent group-level brain responses include some frontal and ventral regions that are not typically observed on individual-level ISC maps. On one hand, this is reminiscent of findings in task-based fMRI that averaging across larger numbers of timeseries “unmasks” the involvement of common task-locked signal in previously unappreciated regions (Gonzalez-Castillo et al., 2012). On the other hand, cross-site similarity between group-level timeseries in some of these regions is, while statistically significant, quite weak, and similar correlations have been interpreted by other groups as showing little evidence of synchronized brain responses (Chang et al., 2021). We see this as a scenario akin to asking “is the glass half empty or half full?”. Weak correlations between time series undoubtedly indicate that the signal is predominantly explained by other sources including endogenous processing, intrinsic brain dynamics, and various sources of scanner and physiological noise. Alternative methods for correcting for multiple comparisons that capture underlying data dimensionality and potential dependencies between timeseries could also shift the statistical threshold delineating which ROIs can be considered weakly correlated above chance. Nonetheless, identifying shared signal – albeit weakly shared – between two datasets is stronger evidence than can be provided by one dataset alone that there is something about these video stimuli that can evoke common brain function in such areas, potentially indirectly and potentially only momentarily. Better understanding the aspects of the video stimulus that drive such weakly evoked responses in brain areas more commonly associated with endogenous brain function is an important topic of future study (see also Chang et al., 2021; Yeshurun et al., 2021).

The specific moments of the video and specific features of the stimulus that drive the most and least consistent brain responses across sites is also a question for further study. A visual comparison across the different video scans presented in Figure 4 shows clear similarities in the patterns of group-level consistency (top) and pairwise across-site ISC (bottom) evoked by all the different video stimuli employed. In other words, brain regions that respond very consistently across sites during one video tend to also respond very consistently in a different video, and vice versa. This surely reflects fundamental aspects of neural architecture for dynamic audiovisual stimulation, as the most consistent brain regions were the primary sensory areas expected to be most directly driven by the stimulus. Some differences in ISC levels across each full-length scan could arise due to differences in video lengths, which varied considerably. But even after equating for video lengths, differences in group-level brain response consistency across different videos could be observed (Supplemental Figure 2). Presumably, these differences are elicited by specific video stimulus features and idiosyncratic responses to those features, as well as the processing demands they impose on the brain (see also Hasson et al., 2010, for discussion of stimulus-specificity of within-site reliability). For instance, cross-site consistency in medial prefrontal cortex (mPFC) for both episodes of *The Office* (Video3 and Video4) appears elevated relative to the other videos. This is noteworthy because *The Office* is a TV show that is characterized by many socially awkward moments and was specifically selected for its increased demands on the social brain (including mPFC; Kennedy & Adolphs, 2012). Further work comprehensively decomposing these videos from low-level stimulus features to high-level semantic properties will be needed to verify this observation and more generally understand how different video stimulus properties influence patterns of consistency across sites.

As noted, the comparison between the unmatched datasets was presented as a case study and as an example with which to contrast the high levels of cross-site similarity in the matched datasets. Particularly with increasing data sharing efforts in recent years, this comparison has more real-world relevance for the pooling of some pre-existing vfMRI datasets, which are unlikely to have been as carefully matched as the primary samples in this study. For the unmatched datasets in the current study, we observed quantitative differences in group-level consistency and pairwise ISC, but qualitatively, the patterns of pairwise ISC remained highly similar across and within each site. For these unmatched datasets, differences in the acquisition and processing varied considerably (Table 1), including participants, scanner model, acquisition parameters including voxel size, sampling rate, multiband parameters, and sequences used for anatomical scans and fieldmaps, and preprocessing choices including denoising methodology, filtering, and smoothing.

Many if not all of these factors could influence cross-site consistency of brain responses (e.g., He et al., 2020; Friedman et al, 2006; Yu et al., 2018). It is also important to note that the levels of consistency observed in the unmatched datasets are not intended to suggest a lower bound. All datasets in this study used the same scanner manufacturer (Siemens) and field strength (3T), and it is reasonable to expect that cross-manufacturer or cross-magnet comparisons could potentially further affect consistency. A full disentangling of the specific combinations of factors that gave rise to the more prevalent differences observed in the unmatched datasets is beyond the scope of the current project, which was not designed to test these factors systematically. An important question for future study would be to unpack these factors by parametrically varying the differences between these datasets, and to include comparisons across different scanner manufacturers and different field strengths. This would also guide the development of statistical harmonization methods for pooling existing video fMRI data (as in Yu et al., 2018; Yamashita et al., 2019, for resting state data), which could span a variety of manufacturers and even field strengths.

Even for the matched datasets, our existing data does not allow us to conclusively separate effects caused by different scanners from other factors that covaried between the matched datasets. Those factors were intentionally minimized, but do include both different physical scanners and different individual subjects. Some aspects of the differences that were observed between these matched datasets could thus have been driven by participant variability rather than scanner differences. To fully decouple individual variability from scanner variability, a new data acquisition with traveling subjects that are repeatedly scanned at different locations (as has been done for resting state designs; Noble et al., 2017) would be required. This would be an important direction for future work.

## Conclusion

In sum, we find similar group-level brain responses spanning the cortex when participants at different sites watch the same video stimulus, and these highly similar average time series occur with both matched and unmatched datasets. When datasets are carefully matched such that the acquisition and processing is effectively identical, differences between datasets at the level of pairwise similarity of individual brain responses are minimal, and some such differences could reflect individual variability rather than scanner-specific effects. When dataset parameters vary more freely, differences between sites are more prevalent, which points to the importance of both careful control for such differences in analyses and of the development of harmonization protocols specific to ISC analyses of video fMRI data for at least some purposes. Nonetheless, the overarching conclusion indicates high levels of consistency in video-evoked fMRI data across these different sites, across matched and unmatched datasets alike. The ability to quantify this consistency highlights one of the unique features of video fMRI and holds promise for further development of this approach to studies of individual differences in healthy and clinical populations alike.

## Acknowledgements

This work was supported in part by the NIH (R01MH110630 and R00MH094409 to DPK and T32HD007475 Postdoctoral Traineeship to LB), the Simons Foundation Autism Research Initiative (RA), and a Della Martin Fellowship (DK). For supercomputing resources, this work was supported in part by Lilly Endowment, Inc., through its support for the Indiana University Pervasive Technology Institute, and in part by the Indiana METACyt Initiative. The Indiana METACyt Initiative at IU was also supported in part by Lilly Endowment, Inc. We thank Susannah Ferguson, Brad Caron, Arispa Weigold, and Steven Lograsso for help with data collection and we are grateful to all our participants and their families.

## Data availability

The primary data (Video1 and Video2) analyzed for this manuscript is publicly in the National Database for Autism Research (NDAR; Hall et al., 2012; https://nda.nih.gov/about.html).

## Author contributions

LB, RA, and DPK conceptualized the project. HC & MT developed MRI protocols, coordinated them across sites, and continuously conducted scanner quality assurance. LB, DK, and Indiana University and Caltech personnel collected data. LB, DK, and YH preprocessed data and ensured data quality. LB & DPK developed the analysis approach and LB analyzed the data. LB drafted the manuscript with input from DPK and all co-authors provided feedback and approved the final version.

