## Supplemental Information for "Video-evoked fMRI BOLD responses are highly consistent across different data acquisition sites"

### Supplemental Methods.

#### Data acquisition, preprocessing, and quality assessment.

An upgrade to the trigger box occurred in the final months of data collection at the IU site, and this sporadically resulted in an intermittent missed trigger and delayed movie start. We used the following algorithm to identify and correct the affected scans. Due to individual variability in hemodynamic response functions, it is expected that under normal circumstances, participant time series in primary sensory areas and the average time series across other subjects in those same regions could be slightly offset. In other words, cross-correlations between participant and average time series would be expected to often peak at a lag of 0, but could also peak at a lag of a small number of TRs. Our goal was to identify scans that were affected by the trigger issue while preserving the individual variability in small offsets that might relate to hemodynamics. Therefore, we began by establishing a baseline in the unaffected data (Caltech data and IU data pre-upgrade, computed separately). Across all movie scans, using the Schaefer400x17 parcellation, we examined 78 subregions from the functional networks containing visual and auditory cortices (bilateral VisCent, VisPeri, and SomMotB networks). For these 78 ROIs, we computed cross-correlations between each participant time series and the median time series across the other participants from the same dataset. We found that in both datasets, the distribution of modal peak lags in the unaffected data was effectively always in the range [-2,2] and with a strong peak at 0 as expected. We therefore considered peak lags <=2 to reflect expected hemodynamic response variability. Next, we computed the same cross-correlations in the data collected after the trigger upgrade. When the modal peak lag across these ROIs exceeded 2, we took this to reflect a delayed movie start, and realigned the BOLD time series by shifting them by the modal peak lag. 35 scans were realigned in total (5 Video1, 9 Video2, 8 Video3, 6 Video4, 3 Video5, 4 Video6). These realignments did not influence the pattern of results reported here, which were effectively identical when conducted with the original (non-realigned) scans.

The following boilerplate text is copied verbatim from the fMRIPrep software reports and is included at the request of the software developers. Note that this text refers to 10 functional scans but only a subset of these scans is included in the current report, and not all the confound time series described were used in our downstream analyses. (For example, we indexed head motion using a different formulation of framewise displacement, see Methods, and did not use DVARs traces, CompCor traces, or measures derived from them in these analyses.)

Results included in this manuscript come from preprocessing performed using fMRIPrep 20.0.7 (Esteban, Markiewicz, et al. (2018); Esteban, Blair, et al. (2018); RRID:SCR_016216), which is based on Nipype 1.4.2 (Gorgolewski et al. (2011); Gorgolewski et al. (2018); RRID:SCR_002502).

*Anatomical data preprocessing*. The T1-weighted (T1w) image was corrected for intensity non-uniformity (INU) with N4BiasFieldCorrection (Tustison et al. 2010), distributed with ANTs 2.2.0 (Avants et al. 2008, RRID:SCR_004757), and used as T1w-reference throughout the workflow. The

T1w-reference was then skull-stripped with a Nipype implementation of the antsBrainExtraction.sh workflow (from ANTs), using OASIS30ANTs as target template. Brain tissue segmentation of cerebrospinal fluid (CSF), white-matter (WM) and gray-matter (GM) was performed on the brain-extracted T1w using fast (FSL 5.0.9, RRID:SCR_002823, Zhang, Brady, and Smith 2001). Brain surfaces were reconstructed using recon-all (FreeSurfer 6.0.1, RRID:SCR_001847, Dale, Fischl, and Sereno 1999), and the brain mask estimated previously was refined with a custom variation of the method to reconcile ANTs-derived and FreeSurfer-derived segmentations of the cortical gray-matter of Mindboggle (RRID:SCR_002438, Klein et al. 2017). Volume-based spatial normalization to one standard space (MNI152Nlin2009cAsym) was performed through nonlinear registration with antsRegistration (ANTs 2.2.0), using brain-extracted versions of both T1w reference and the T1w template. The following template was selected for spatial normalization: ICBM 152 Nonlinear Asymmetrical template version 2009c [Fonov et al. (2009), RRID:SCR_008796; TemplateFlow ID: MNI152Nlin2009cAsym].

*Functional data preprocessing*. For each of the 10 BOLD runs found per subject (across all tasks and sessions), the following preprocessing was performed. First, a reference volume and its skull-stripped version were generated using a custom methodology of fMRIPrep. A B0-nonuniformity map (or fieldmap) was estimated based on two (or more) echo-planar imaging (EPI) references with opposing phase-encoding directions, with 3dQwarp Cox and Hyde (1997) (AFNI 20160207). Based on the estimated susceptibility distortion, a corrected EPI (echo-planar imaging) reference was calculated for a more accurate co-registration with the anatomical reference. The BOLD reference was then co-registered to the T1w reference using bbregister (FreeSurfer) which implements boundary-based registration (Greve and Fischl 2009). Co-registration was configured with six degrees of freedom. Head-motion parameters with respect to the BOLD reference (transformation matrices, and six corresponding rotation and translation parameters) are estimated before any spatiotemporal filtering using mcflirt (FSL 5.0.9, Jenkinson et al. 2002). The BOLD time-series were resampled onto the following surfaces (FreeSurfer reconstruction nomenclature): fsaverage5, fsaverage6, fsnative. The BOLD time-series (including slice-timing correction when applied) were resampled onto their original, native space by applying a single, composite transform to correct for head-motion and susceptibility distortions. These resampled BOLD time-series will be referred to as preprocessed BOLD in original space, or just preprocessed BOLD. The BOLD time-series were resampled into standard space, generating a preprocessed BOLD run in MNI152Nlin2009cAsym space. First, a reference volume and its skull-stripped version were generated using a custom methodology of fMRIPrep. Several confounding time-series were calculated based on the preprocessed BOLD: framewise displacement (FD), DVARS and three region-wise global signals. FD and DVARS are calculated for each functional run, both using their implementations in Nipype (following the definitions by Power et al. 2014). The three global signals are extracted within the CSF, the WM, and the whole-brain masks. Additionally, a set of physiological regressors were extracted to allow for component-based noise correction (CompCor, Behzadi et al. 2007). Principal components are estimated after high-pass filtering the preprocessed BOLD time-series (using a discrete cosine filter with 128s cut-off) for the two CompCor variants: temporal (tCompCor) and anatomical (aCompCor). tCompCor components are then calculated from the top 5% variable voxels within a mask covering the subcortical regions. This subcortical mask is obtained by heavily eroding the brain mask, which ensures it does not include cortical GM regions. For aCompCor, components are calculated within the intersection of the aforementioned mask and the union of CSF and WM masks calculated in T1w space, after their projection to the native space of each functional run (using the inverse BOLD-to-T1w transformation). Components are also calculated separately within the WM and CSF masks. For each CompCor decomposition, the k components with the largest singular values are retained, such that the retained components' time series are sufficient to explain 50 percent of variance across the nuisance mask (CSF, WM, combined, or temporal). The remaining components are dropped from consideration. The head-motion estimates calculated in the correction step were also placed within the corresponding confounds file. The confound time series derived from head motion estimates and global signals were expanded with the inclusion of temporal derivatives and quadratic terms for each (Satterthwaite et al. 2013). Frames that exceeded a threshold of 0.4 mm FD or 1.5 standardised DVARS were annotated as motion outliers. All resamplings can be performed with a single interpolation step by composing all the pertinent transformations (i.e. head-motion transform matrices, susceptibility distortion correction when available, and co-registrations to anatomical and output spaces). Gridded (volumetric) resamplings were performed using antsApplyTransforms (ANTs), configured with Lanczos interpolation to minimize the smoothing effects of other kernels (Lanczos 1964). Non-gridded (surface) resamplings were performed using mri_vol2surf (FreeSurfer).

Many internal operations of fMRIPrep use Nilearn 0.6.2 (Abraham et al. 2014, RRID:SCR_001362), mostly within the functional processing workflow. For more details of the pipeline, see the section corresponding to workflows in fMRIPrep's documentation.

### Supplemental Figures

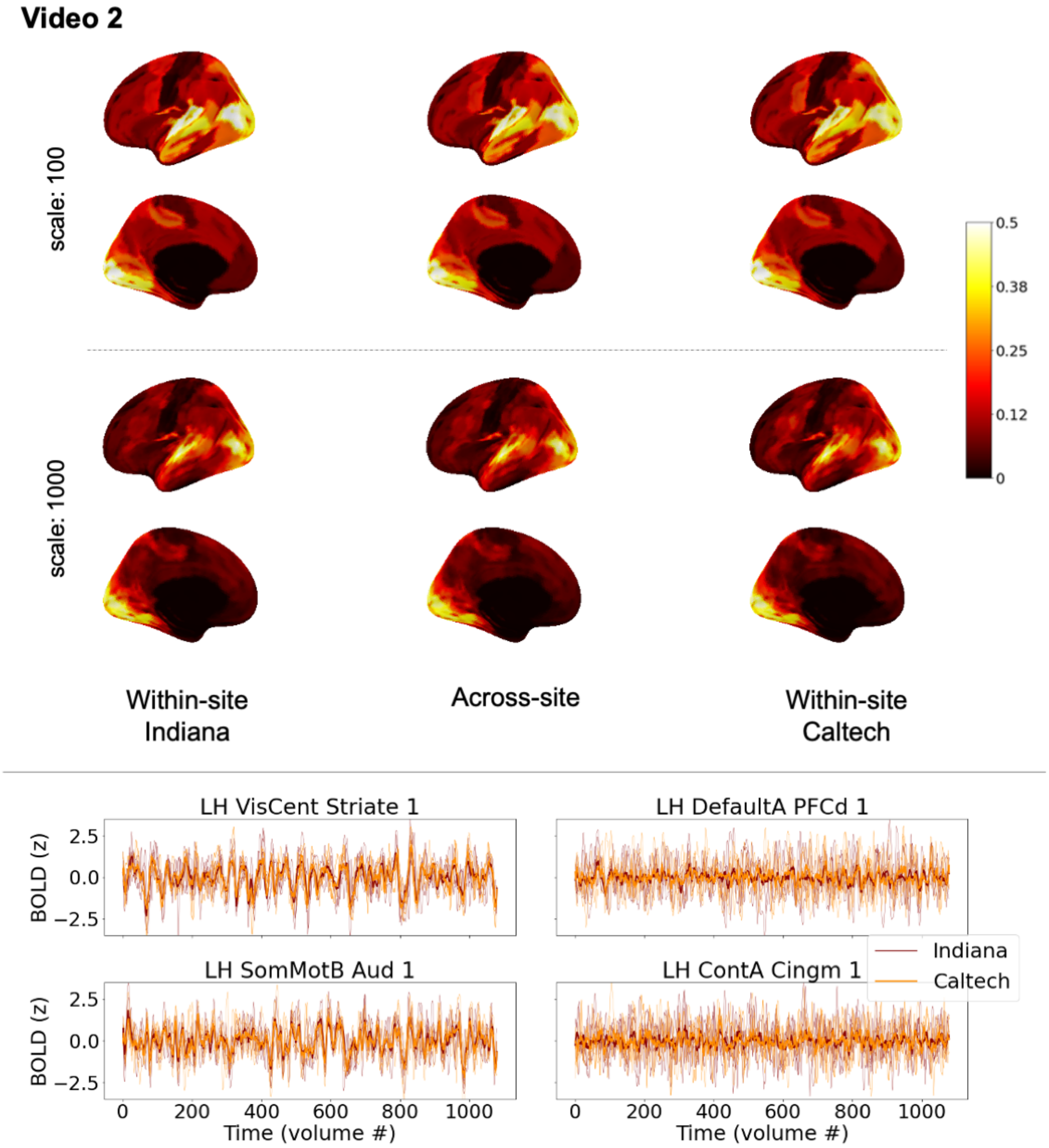


*Supplemental Figure 1. Similarity of individual participant brain responses within and across sites during Video2, for the matched datasets. Top: brain maps depict magnitudes of medians of pairwise correlations between participant brain response time series in each region, for the most coarse (top) and the most fine (bottom) scales of the Schaefer cortical parcellation. The left and right columns show*

*correlations among pairs of participants at the same site (left: Indiana; right: Caltech). The center column shows correlations among pairs of participants spanning different sites. While absolute values are depicted here for readability, nearly all median correlations were positive, except seven ROIs with effectively zero median correlations above -0.001. As in Figure 2, black along the midline in medial views indicates missing data, not low correlations. See also Figure 4 (bottom panel, second plot) for a line version of this same data, and Supplemental Table 2 for characterization of differences and effect sizes. Bottom: line plots depict timeseries from 5 randomly-selected participants at each site in primary sensory areas (left) and association areas (right), along with the site-level median time series shown in Figure 2. These line plots use the same mid-scale parcellation as Figure 2 (Schaefer 400x17). See also Figure 3 for the equivalent figure for Video1.*

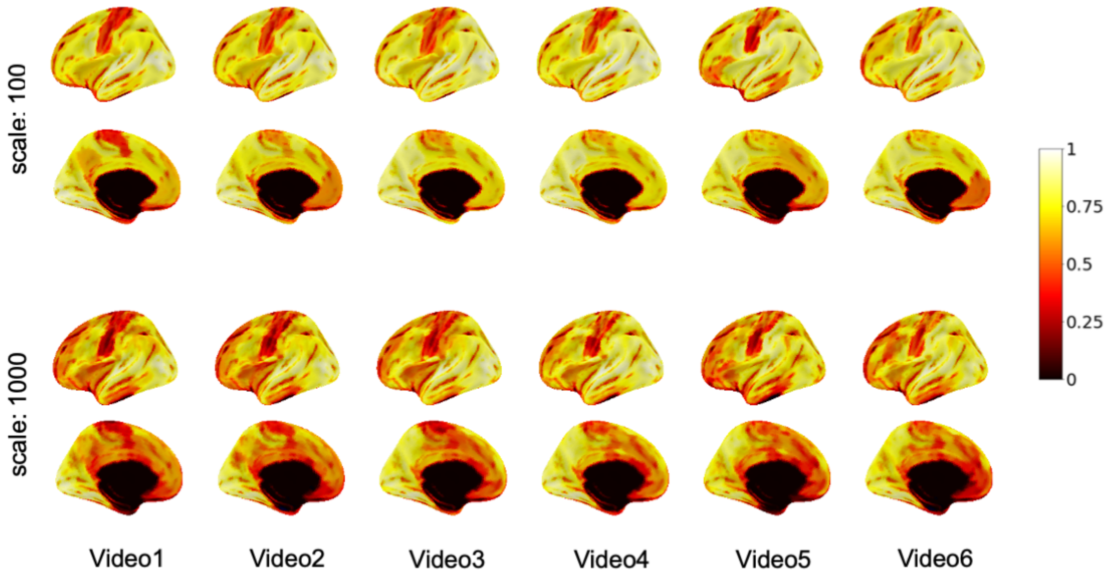


*Supplemental Figure 2. Randomly downsampled version of figure showing consistency in brain responses across sites while participants watched the same video, across a variety of different videos, in the matched datasets. To account for differences in scan length, contiguous segments of 232 TRs (half the length of the shortest scan) were randomly selected from each scan, and the correlation between the*

*group average timeseries for each site during that segment was computed for each ROI. This figure presents in each ROI the median of these correlations across 100 randomly selected segments. Patterns are largely the same as seen in Figure 4 (top), which preserves the entire scan length. Only parcellation scales of 100 and 1000 ROIs are shown for space considerations. As in Figure 2, black along the midline in medial views indicates missing data, not low correlations.*

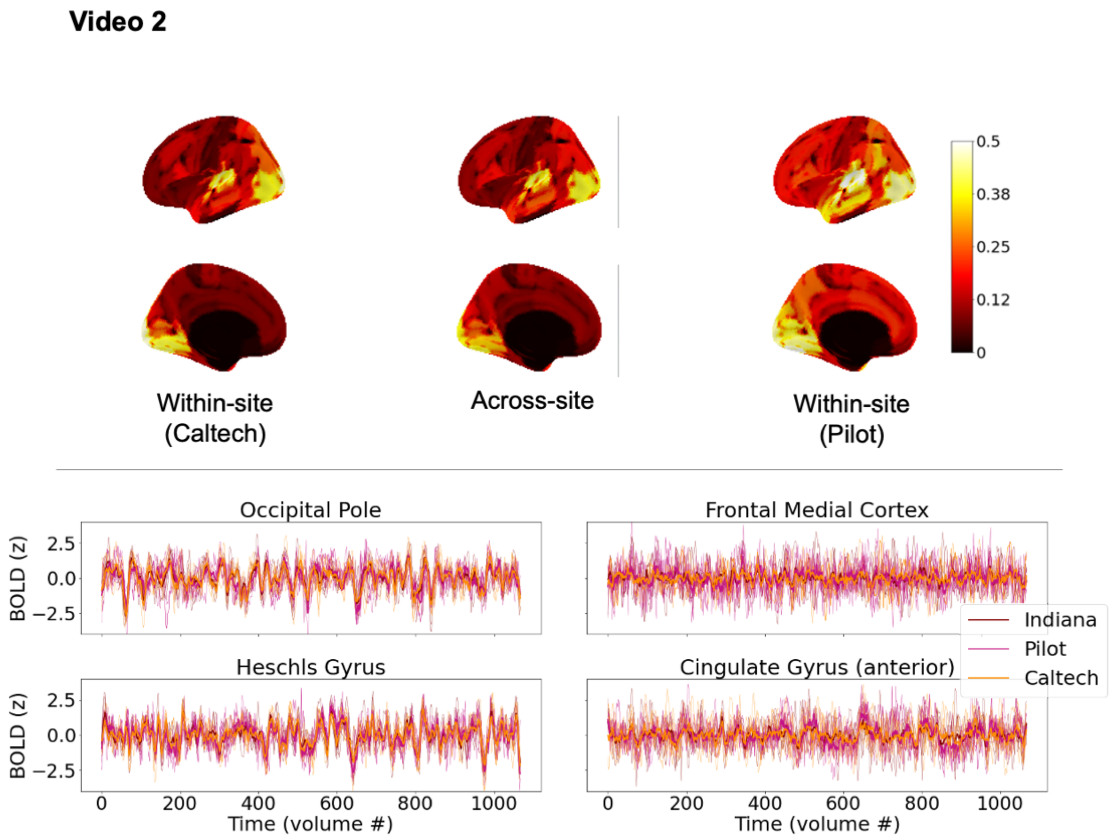


*Supplemental Figure 3. Exploratory comparison of brain response consistency across unmatched datasets, at the level of individual time series (Fig. 1B), for Video2. Top: brain maps depict magnitudes of medians of pairwise correlations between participant brain response time series in each region. The left and right columns show correlations among pairs of participants at the same site (left: Caltech; right: Pilot). The center column shows correlations among pairs of participants spanning different sites (as in Fig. 1B, green). Black along the midline in medial views indicates missing data, not low correlations. Center: Time series plots show five randomly-selected individual time series for each site in*

*sensory areas (left) and association areas (right), along with the median time series across all participants at that site superimposed in bold. Bottom: Within- and across-site ISC values from top panel (Video1) and Supplemental Figure 3 (Video2) presented as a line plot, to facilitate comparison. See text for summary of differences and effect sizes. As in Figure 4 (bottom), individual data points are connected with a line, but these plots are not time series. Rather, each data point reflects median similarity across pairs of timeseries. When values for a given ROI are similar across the two different scans, that reflects comparable levels of similarity across entirely different brain response time series evoked by different videos. All panels use the Harvard-Oxford 96-ROI cortical parcellation. See also Figure 6 for the equivalent figure for Video1.*

### Supplemental Tables

Supplemental Table 1.

*Key to x tick labels (ROI names) in Figures 4 and 6.*

| X value (ROI #) | ROI name in Fig 4 (center) (Schaefer100x17) | ROI name in Fig 6 (Harvard-Oxford cortical) |
|---|---|---|
| 0 | LH_VisCent_ExStr_1 | LH_FrontalPole |
| 1 | LH_VisCent_ExStr_2 | LH_InsularCortex |
| 2 | LH_VisCent_Striate_1 | LH_SuperiorFrontalGyrus |
| 3 | LH_VisCent_ExStr_3 | LH_MiddleFrontalGyrus |
| 4 | LH_VisPeri_ExStrInf_1 | LH_InferiorFrontalGyrusparstriangularis |
| 5 | LH_VisPeri_StriCal_1 | LH_InferiorFrontalGyrusparsopercularis |
| 6 | LH_VisPeri_ExStrSup_1 | LH_PrecentralGyrus |
| 7 | LH_SomMotA_1 | LH_TemporalPole |
| 8 | LH_SomMotA_2 | LH_SuperiorTemporalGyrusanteriordivision |
| 9 | LH_SomMotB_Aud_1 | LH_SuperiorTemporalGyrusposteriordivision |
| 10 | LH_SomMotB_S2_1 | LH_MiddleTemporalGyrusanteriordivision |
| 11 | LH_SomMotB_S2_2 | LH_MiddleTemporalGyrusposteriordivision |
| 12 | LH_SomMotB_Cent_1 | LH_MiddleTemporalGyrustemporooccipitalpart |
| 13 | LH_DorsAttnA_TempOcc_1 | LH_InferiorTemporalGyrusanteriordivision |
| 14 | LH_DorsAttnA_ParOcc_1 | LH_InferiorTemporalGyrusposteriordivision |

| | | |
|---|---|---|
| 15 | LH_DorsAttnA_SPL_1 | LH_InferiorTemporalGyrustemporooccipitalpart |
| 16 | LH_DorsAttnB_PostC_1 | LH_PostcentralGyrus |
| 17 | LH_DorsAttnB_PostC_2 | LH_SuperiorParietalLobule |
| 18 | LH_DorsAttnB_PostC_3 | LH_SupramarginalGyrusanteriordivision |
| 19 | LH_DorsAttnB_FEF_1 | LH_SupramarginalGyrusposteriordivision |
| 20 | LH_SalVentAttnA_ParOper_1 | LH_AngularGyrus |
| 21 | LH_SalVentAttnA_Ins_1 | LH_LateralOccipitalCortexsuperiordivision |
| 22 | LH_SalVentAttnA_Ins_2 | LH_LateralOccipitalCortexinferiordivision |
| 23 | LH_SalVentAttnA_ParMed_1 | LH_IntracalcarineCortex |
| 24 | LH_SalVentAttnA_FrMed_1 | LH_FrontalMedialCortex |
| 25 | LH_SalVentAttnB_PFCl_1 | LH_JuxtapositionalLobuleCortex |
| 26 | LH_SalVentAttnB_PFCmp_1 | LH_SubcallosalCortex |
| 27 | LH_LimbicB_OFC_1 | LH_ParacingulateGyrus |
| 28 | LH_LimbicA_TempPole_1 | LH_CingulateGyrusanteriordivision |
| 29 | LH_LimbicA_TempPole_2 | LH_CingulateGyrusposteriordivision |
| 30 | LH_ContA_IPS_1 | LH_PrecuneousCortex |
| 31 | LH_ContA_PFCl_1 | LH_CunealCortex |
| 32 | LH_ContA_PFCl_2 | LH_FrontalOrbitalCortex |
| 33 | LH_ContB_PFClv_1 | LH_ParahippocampalGyrusanteriordivision |
| 34 | LH_ContC_pCun_1 | LH_ParahippocampalGyrusposteriordivision |
| 35 | LH_ContC_pCun_2 | LH_LingualGyrus |
| 36 | LH_ContC_Cingp_1 | LH_TemporalFusiformCortexanteriordivision |
| 37 | LH_DefaultA_PFCd_1 | LH_TemporalFusiformCortexposteriordivision |
| 38 | LH_DefaultA_pCunPCC_1 | LH_TemporalOccipitalFusiformCortex |
| 39 | LH_DefaultA_PFCm_1 | LH_OccipitalFusiformGyrus |
| 40 | LH_DefaultB_Temp_1 | LH_FrontalOperculumCortex |
| 41 | LH_DefaultB_Temp_2 | LH_CentralOpercularCortex |
| 42 | LH_DefaultB_IPL_1 | LH_ParietalOperculumCortex |
| 43 | LH_DefaultB_PFCd_1 | LH_PlanumPolare |
| 44 | LH_DefaultB_PFCl_1 | LH_HeschlsGyrus |
| 45 | LH_DefaultB_PFCv_1 | LH_PlanumTemporale |
| 46 | LH_DefaultB_PFCv_2 | LH_SupracalcarineCortex |
| 47 | LH_DefaultC_Rsp_1 | LH_OccipitalPole |
| 48 | LH_DefaultC_PHC_1 | RH_FrontalPole |
| 49 | LH_TempPar_1 | RH_InsularCortex |
| 50 | RH_VisCent_ExStr_1 | RH_SuperiorFrontalGyrus |
| 51 | RH_VisCent_ExStr_2 | RH_MiddleFrontalGyrus |
| 52 | RH_VisCent_ExStr_3 | RH_InferiorFrontalGyrusparstriangularis |
| 53 | RH_VisPeri_StriCal_1 | RH_InferiorFrontalGyrusparsopercularis |

| | | |
|---|---|---|
| 54 | RH_VisPeri_ExStrInf_1 | RH_PrecentralGyrus |
| 55 | RH_VisPeri_ExStrSup_1 | RH_TemporalPole |
| 56 | RH_SomMotA_1 | RH_SuperiorTemporalGyrusanteriordivision |
| 57 | RH_SomMotA_2 | RH_SuperiorTemporalGyrusposteriordivision |
| 58 | RH_SomMotA_3 | RH_MiddleTemporalGyrusanteriordivision |
| 59 | RH_SomMotA_4 | RH_MiddleTemporalGyrusposteriordivision |
| 60 | RH_SomMotB_Aud_1 | RH_MiddleTemporalGyrustemporooccipitalpart |
| 61 | RH_SomMotB_S2_1 | RH_InferiorTemporalGyrusanteriordivision |
| 62 | RH_SomMotB_S2_2 | RH_InferiorTemporalGyrusposteriordivision |
| 63 | RH_SomMotB_Cent_1 | RH_InferiorTemporalGyrustemporooccipitalpart |
| 64 | RH_DorsAttnA_TempOcc_1 | RH_PostcentralGyrus |
| 65 | RH_DorsAttnA_ParOcc_1 | RH_SuperiorParietalLobule |
| 66 | RH_DorsAttnA_SPL_1 | RH_SupramarginalGyrusanteriordivision |
| 67 | RH_DorsAttnB_PostC_1 | RH_SupramarginalGyrusposteriordivision |
| 68 | RH_DorsAttnB_PostC_2 | RH_AngularGyrus |
| 69 | RH_DorsAttnB_FEF_1 | RH_LateralOccipitalCortexsuperiordivision |
| 70 | RH_SalVentAttnA_ParOper_1 | RH_LateralOccipitalCortexinferiordivision |
| 71 | RH_SalVentAttnA_Ins_1 | RH_IntracalcarineCortex |
| 72 | RH_SalVentAttnA_ParMed_1 | RH_FrontalMedialCortex |
| 73 | RH_SalVentAttnA_FrMed_1 | RH_JuxtapositionalLobuleCortex |
| 74 | RH_SalVentAttnB_IPL_1 | RH_SubcallosalCortex |
| 75 | RH_SalVentAttnB_PFCl_1 | RH_ParacingulateGyrus |
| 76 | RH_SalVentAttnB_PFCmp_1 | RH_CingulateGyrusanteriordivision |
| 77 | RH_LimbicB_OFC_1 | RH_CingulateGyrusposteriordivision |
| 78 | RH_LimbicA_TempPole_1 | RH_PrecuneousCortex |
| 79 | RH_ContA_IPS_1 | RH_CunealCortex |
| 80 | RH_ContA_PFCl_1 | RH_FrontalOrbitalCortex |
| 81 | RH_ContA_PFCl_2 | RH_ParahippocampalGyrusanteriordivision |
| 82 | RH_ContB_Temp_1 | RH_ParahippocampalGyrusposteriordivision |
| 83 | RH_ContB_IPL_1 | RH_LingualGyrus |
| 84 | RH_ContB_PFCld_1 | RH_TemporalFusiformCortexanteriordivision |
| 85 | RH_ContB_PFClv_1 | RH_TemporalFusiformCortexposteriordivision |
| 86 | RH_ContC_Cingp_1 | RH_TemporalOccipitalFusiformCortex |
| 87 | RH_ContC_pCun_1 | RH_OccipitalFusiformGyrus |
| 88 | RH_DefaultA_IPL_1 | RH_FrontalOperculumCortex |
| 89 | RH_DefaultA_PFCd_1 | RH_CentralOpercularCortex |
| 90 | RH_DefaultA_pCunPCC_1 | RH_ParietalOperculumCortex |
| 91 | RH_DefaultA_PFCm_1 | RH_PlanumPolare |
| 92 | RH_DefaultB_PFCd_1 | RH_HeschlsGyrus |

| | | |
|---|---|---|
| 93 | RH_DefaultB_PFCv_1 | RH_PlanumTemporale |
| 94 | RH_DefaultB_PFCv_2 | RH_SupracalcarineCortex |
| 95 | RH_DefaultC_Rsp_1 | RH_OccipitalPole |
| 96 | RH_DefaultC_PHC_1 | - |
| 97 | RH_TempPar_1 | - |
| 98 | RH_TempPar_2 | - |
| 99 | RH_TempPar_3 | - |

Note: Table presents the ROI labels corresponding to the *x* axis ticks in the line plots for Figures 4 (100-scale parcellation) and 6. Schaefer100x17 labels reflect the ordering returned by the fetch_atlas_schaefer_2018 routine in the nilearn package and are reprinted here for convenience. Please consult fetch_atlas_schaefer_2018 for the Schaefer1000x17 labels corresponding to Figure 4 (1000-scale parcellation), which are too long to print. Tick labels are omitted from the figure for readability.

Supplemental Table 2.

*Differences and effect sizes for pairwise within-site versus across-site ISC comparisons.*

| | Video1 | | Video2 | | Video3 | | Video4 | | Video5 | | Video6 | |
|---|---|---|---|---|---|---|---|---|---|---|---|---|
| scale | *IU* | *Cal* | *IU* | *Cal* | *IU* | *Cal* | *IU* | *Cal* | *IU* | *Cal* | *IU* | *Cal* |
| 100 | 0.005 (0.01) | -0.001 (0.02) | 0.006 (0.02) | 0.001 (0.02) | 0.003 (0.01) | 0.0* (0.01) | 0.0* (0.02) | 0.003 (0.02) | 0.02 (0.02) | -0.01 (0.02) | 0.02 (0.02) | -0.005 (0.02) |
| | 0.51 (0.04) | 0.49 (0.04) | 0.51 (0.04) | 0.48 (0.04) | 0.5 (0.04) | 0.48 (0.05) | 0.49 (0.05) | 0.49 (0.05) | 0.53 (0.03) | 0.45 (0.04) | 0.55 (0.05) | 0.47 (0.05) |
| 200 | 0.004 (0.01) | -0.0* (0.02) | 0.005 (0.01) | -0.0* (0.02) | 0.003 (0.01) | -0.002 (0.01) | 0.002 (0.01) | -0.0* (0.02) | 0.01 (0.02) | -0.009 (0.02 | 0.02 (0.02) | -0.004 (0.02) |
| | 0.51 (0.04) | 0.49 (0.04) | 0.51 (0.04) | 0.48 (0.04) | 0.5 (0.04) | 0.48 (0.04) | 0.5 (0.04) | 0.49 (0.04) | 0.52 (0.04) | 0.47 (0.04) | 0.53 (0.05) | 0.47 (0.04) |
| 400 | 0.004 (0.01) | 0.002 (0.02) | 0.005 (0.01) | 0.0* (0.01) | 0.003 (0.01) | -0.0* (0.02) | 0.0* (0.01) | 0.002 (0.02) | 0.01 (0.02) | -0.008 (0.02) | 0.02 (0.02) | -0.004 (0.02) |
| | 0.51 (0.04) | 0.49 (0.04) | 0.51 (0.04) | 0.48 (0.04) | 0.5 (0.04) | 0.48 (0.04) | 0.5 (0.04) | 0.49 (0.05) | 0.52 (0.04) | 0.47 (0.05) | 0.54 (0.05) | 0.47 (0.05) |
| 600 | 0.004 (0.01) | -0.0* (0.02) | 0.005 (0.01) | -0.0* (0.02) | 0.004 (0.01) | -0.002 (0.02) | 0.001 (0.01) | 0.001 (0.02) | 0.01 (0.02) | -0.009 (0.02) | 0.02 (0.02) | -0.004 (0.02) |
| | 0.51 (0.04) | 0.49 (0.04) | 0.51 (0.04) | 0.48 (0.04) | 0.5 (0.04) | 0.48 (0.04) | 0.49 (0.04) | 0.49 (0.05) | 0.52 (0.04) | 0.47 (0.04) | 0.53 (0.05) | 0.47 (0.05) |
| 800 | 0.004 (0.01) | -0.0* (0.02) | 0.04 (0.01) | -0.001 (0.02) | 0.004 (0.01) | -0.002 (0.01) | 0.002 (0.01) | 0.001 (0.02) | 0.01 (0.02) | -0.009 (0.02) | 0.02 (0.02) | -0.004 (0.02) |
| | 0.51 (0.04) | 0.49 (0.04 | 0.51 (0.04) | 0.48 (0.04 | 0.5 (0.04) | 0.48 (0.04) | 0.5 (0.04) | 0.49 (0.04) | 0.52 (0.04) | 0.47 (0.04) | 0.53 (0.05) | 0.48 (0.05) |

| | | | | | | | | | | | | |
|---|---|---|---|---|---|---|---|---|---|---|---|---|
| 1000 | 0.004 (0.01) | 0.0* (0.02) | 0.005 (0.01) | -0.0* (0.02 | 0.003 (0.01) | -0.01 (0.01) | 0.002 (0.01) | 0.0* (0.02) | 0.01 (0.02) | -0.008 (0.02) | 0.02 (0.02) | -0.004 (0.02) |
| | 0.51 (0.04) | 0.49 (0.04) | 0.51 (0.04) | 0.48 (0.04) | 0.5 (0.04) | 0.48 (0.04) | 0.5 (0.04) | 0.49 (0.04) | 0.52 (0.04) | 0.47 (0.04) | 0.53 (0.05) | 0.48 (0.05) |

Note: Site labels (IU, Cal) refers to comparisons between within-site ISC at that site and across-site ISC. Values are presented as follows. For each parcellation scale, the first row shows the median (IQR) of the differences between the median within-site ISC and the median across-site ISC, across all ROIs in the parcellation. The second row (in gray) shows the median (IQR) of the common-language effect sizes (CLES) comparing the distributions of within-site ISC and across-site ISC, across all ROIs in the parcellation. For all comparisons, the direction is within – across: positive differences, and effect sizes above 0.5, indicate larger within-site than across-site ISC (and vice versa). Very small values between +/- 0.001 are truncated to +/- 0.0 for space and marked with an asterisk.

Jenkinson, Mark, Peter Bannister, Michael Brady, and Stephen Smith. 2002. "Improved Optimization for the Robust and Accurate Linear Registration and

Motion Correction of Brain Images." NeuroImage 17 (2): 825–41. https://doi.org/10.1006/nimg.2002.1132.

Klein, Arno, Satrajit S. Ghosh, Forrest S. Bao, Joachim Giard, Yrjö Häme, Eliezer Stavsky, Noah Lee, et al. 2017. "Mindboggling Morphometry of Human Brains." PLOS Computational Biology 13 (2): e1005350. https://doi.org/10.1371/journal.pcbi.1005350.

Lanczos, C. 1964. "Evaluation of Noisy Data." Journal of the Society for Industrial and Applied Mathematics Series B Numerical Analysis 1 (1): 76–85. https://doi.org/10.1137/0701007.

Power, Jonathan D., Anish Mitra, Timothy O. Laumann, Abraham Z. Snyder, Bradley L. Schlaggar, and Steven E. Petersen. 2014. "Methods to Detect, Characterize, and Remove Motion Artifact in Resting State fMRI." NeuroImage 84 (Supplement C): 320–41. https://doi.org/10.1016/j.neuroimage.2013.08.048.

Satterthwaite, Theodore D., Mark A. Elliott, Raphael T. Gerraty, Kosha Ruparel, James Loughead, Monica E. Calkins, Simon B. Eickhoff, et al. 2013. "An improved framework for confound regression and filtering for control of motion artifact in the preprocessing of resting-state functional connectivity data." NeuroImage 64 (1): 240–56. https://doi.org/10.1016/j.neuroimage.2012.08.052.

Tustison, N. J., B. B. Avants, P. A. Cook, Y. Zheng, A. Egan, P. A. Yushkevich, and J. C. Gee. 2010. "N4ITK: Improved N3 Bias Correction." IEEE Transactions on Medical Imaging 29 (6): 1310–20. https://doi.org/10.1109/TMI.2010.2046908.

Zhang, Y., M. Brady, and S. Smith. 2001. "Segmentation of Brain MR Images Through a Hidden Markov Random Field Model and the Expectation-Maximization Algorithm." IEEE Transactions on Medical Imaging 20 (1): 45–57. https://doi.org/10.1109/42.906424.
